# Multiscale wrinkling dynamics in epithelial shells

**DOI:** 10.1101/2025.06.30.662426

**Authors:** Nimesh Chahare, Adam Ouzeri, Thomas Wilson, Pradeep K. Bal, Tom Golde, Guillermo Vilanova, Pau Pujol-Vives, Pere Roca-Cusachs, Xavier Trepat, Marino Arroyo

**Author notes:** Correspondence to: Xavier Trepat, PhD, Institute for Bioengineering of Catalonia, Barcelona, Spain, Marino Arroyo, PhD, Universitat Politècnica de Catalunya, Barcelona, Spain.

## Abstract

Thin shells buckle and wrinkle when compressed. While this behavior is generally detrimental in engineering, it has been widely implicated in epithelial morphogenesis and patterning during development. Yet the rules governing buckling of active viscoelastic shells like epithelia remain unclear. Here we delineate those rules by combining an experimental system that allows us to sculpt epithelial shells and subject them to controlled deflation with a 3D computational model linking cytoskeletal dynamics to tissue mechanics. Experiments and simulations across several orders of magnitude in time and space reveal that buckling emerges for fast deflation relative to the cortex’s relaxation time, and is suppressed by high contractility. We show, further, that the tissue develops wrinkle patterns with different degrees of symmetry breaking that depend on its size and viscous confinement. Strikingly, we find that epithelial buckling is a multiscale phenomenon involving long-lived supracellular folds but also short-lived subcellular wrinkles in the actin cortex. Finally, by forming epithelial shells with anisotropic curvature we rationally direct buckling into predictable wrinkle patterns. Our study shows that epithelial tissues can be understood as hierarchical materials with mechanical instabilities that can be harnessed to engineer epithelial morphogenesis.

## Main Text

Thin-shell structures are ubiquitous both in natural and engineered systems, ranging from microscale carbon nanotubes^1^ and viral capsids^2^ to colossal dome-shaped buildings and lithospheric slabs^3^. It is an everyday experience that slender shells buckle under compressive stresses, such as when we squash a soda can, poke a beach ball or bend a plastic straw. These buckling-induced morphological instabilities provide a mechanism to relax in-plane elastic energy at the expense of energetically less costly bending^4^. Buckling of thin shells generates large deflections and can lead to structural failure^5^, but it can also produce wrinkling patterns depending on shell geometry, mechanical properties, confinement, and applied loads^4,6^. These wrinkling patterns can, in turn, be harnessed to engineer surfaces with tunable optical, adhesive, or aerodynamic properties^7–9^.

In biology, surface patterns arising from buckling instabilities have been shown to control morphogenetic processes in the skin, brain^10^ or gut of animals^11^, in leaves and flowers^12^, and in bacterial biofilms^13^. In epithelial tissues lining organs such as the gut, lung and skin, buckling instabilities have been involved in the formation and maintenance of functional patterned folds such as villi or alveoli^14^. However, unlike inert shells, the mechanisms by which epithelial shells buckle to generate patterned folds and wrinkles are poorly understood, both experimentally and theoretically. This challenge arises from the distinctive mechanical features of epithelial sheets, which include the hierarchical organization of stress bearing elements, the stress-buffering mechanisms operating across different length and time scales^15^, and the presence of active surface tension generated by the contractile actomyosin cortex^16^. In addition, we lack experimental tools to induce buckling of curved epithelia under controlled mechanical and geometrical conditions, as well as theoretical and computational models capturing their hierarchical organization and active viscoelastic mechanics.

Here we establish the multiscale buckling dynamics of an epithelial monolayer as a function of its main mechanical and geometrical determinants. Inspired by deflation assays commonly used to study buckling in inert spherical shells^6,17–19^, we introduce a new approach to form epithelial shells and then deflate them to induce buckling instabilities. This approach is based on a microfluidic chip that allows us to form and manipulate an epithelial shell of controlled geometry. Our findings reveal that tissue buckling is a multiscale phenomenon, characterized by wrinkling patterns with dynamics that depend non-trivially on tissue slenderness, active surface tension, and the time scales of mechanical manipulation. We also present a 3D model of the epithelium, showing that the observed phenomenology can be largely explained by the active viscoelastic properties of the actomyosin cortex. Lastly, we demonstrate how these active viscoelastic properties, when combined with adhesion micropatterning, can be leveraged to engineer epithelial wrinkles with predictable geometries.

### Engineering epithelial buckling

We designed a microfluidic chip called “MOLI” (Mono-Layer Inflator), in which a Madin-Darby canine kidney (MDCK) cell monolayer is seeded on a porous membrane coated with patterned regions of high and low concentration of the adhesive protein fibronectin (Fig. 1a, Extended Fig. 1). By applying hydrostatic pressure across this membrane, the cell monolayer selectively delaminates in the region of low adhesion (the footprint) while remaining attached in the region of high-adhesion. This results in a suspended curved monolayer (referred to hereafter as shell) whose shape and size can be controlled by the applied pressure and fibronectin pattern, and whose dynamics can be followed by confocal or light-sheet microscopy (Fig. 1a,b, Extended Fig. 1). Upon a sudden pressure increase of 200 Pa, cell monolayers delaminated from the low-adherent patterns and reached a steady-state shape and tension within a few minutes (Fig. 1c, Extended Fig. 2). Plateau tissue tensions were found in the 5 to 10 mN/m range, commensurate with active tensions produced by the actomyosin cortex^20,21^. This behavior is qualitatively and quantitatively consistent with the current view that, within a relatively large range of stretch and in the absence of neighbor exchanges, epithelial mechanical properties are dominated by the sub-cellular dynamics of the actomyosin cytoskeleton^20,22,23^. This active gel lining the plasma membrane of epithelial cells can be described as a contractile viscoelastic fluid layer made of filaments, crosslinkers, and myosin motors, all of which undergo turnover (Fig. 1d). At short times, it behaves as an elastic pre-stressed network, but over a viscoelastic relaxation time *τ*_ve_ of tens of seconds to minutes, elastic tension dissipates and only active tension remains^24,25^ (Fig. 1e).

**Figure 1:**
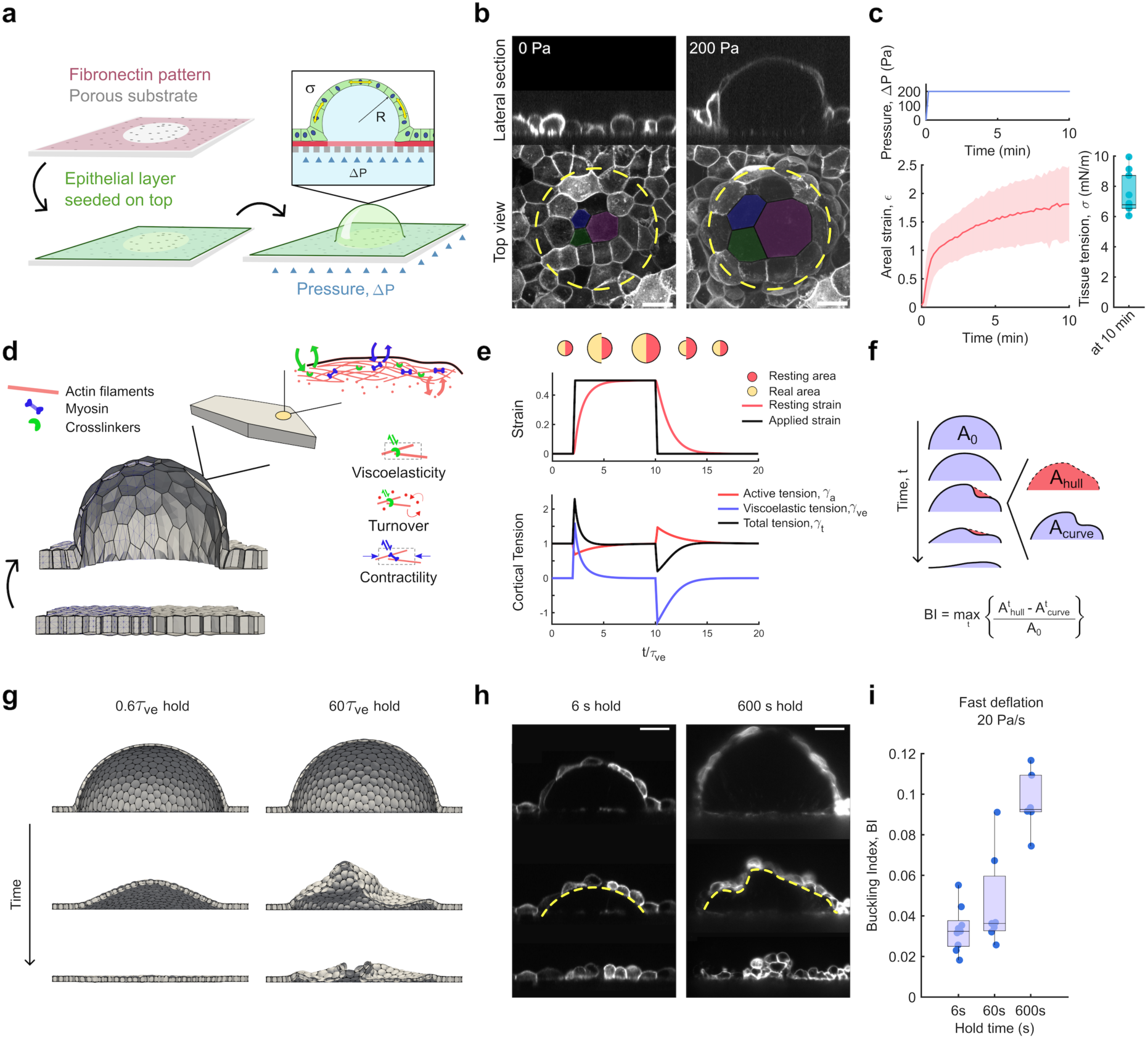
Engineering epithelial shell buckling. **a**, Scheme of the experimental setup. A porous substrate is patterned with high and low fibronectin regions (pink). MDCK cells are seeded on the substrate and allowed to reach confluence to form a tight epithelial monolayer (green). Upon applying positive pressure (blue), the epithelial monolayer delaminates from the low fibronectin areas and inflates to form a shell. **b**, Representative confocal images of the epithelial monolayer at 0 Pa and 200 Pa pressure. A dashed line represents the shell footprint and colored cells illustrate the extent of cell stretching. Top: lateral view. Bottom: maximum intensity projection (Scale bar, 20 µm). **c**, Time evolution of the shell’s areal strain *ε* (bottom left) and tension *σ* after 10 min (bottom right) in response to a pressure step Δ*P* of 200 Pa (top). Data shown as mean ± SD (shaded area) of n = 10 shells. **d**, Scheme of our 3D active gel vertex model (see Supplementary Note 1). The tissue is modeled as an ensemble of triangulated active cortical surfaces enclosing constant cellular volumes. Each surface exhibits viscoelasticity, turnover dynamics, and active contractility representative of the actomyosin network,. **e,** Behavior of the active gel that forms each cortical surface. Following a step stretch, viscoelastic tension increases and then gradually relaxes as the resting strain increases towards the actual strain, whereas the active tension slightly decreases due to cortical dilution and recovers due to turnover. The opposite trends operate upon unstretch. **f,** Scheme representing shell buckling quantification by tracking the shell’s midsection. The Buckling Index (*BI*) is the maximum difference between the area under the curve (*A*_*curve*_) and its convex hull (*A*_ℎ*ull*_), relative to the maximum value of *A*_*curve*_ (*A*_0_). **g,** Simulations of rapid deflation events following short (left) and fast (right) hold times compared to the viscoelastic relaxation time (Movie 1). Estimating *τ*_ve_ ≈ 10 s, the hold times are 6 s and 600 s. **h,** Representative light sheet microscope images of shell mid-sections under the same conditions as in **g**. Yellow dashed lines highlight shape of the monolayer in these two conditions. Scale bar, 20 µm. **i,** The buckling index of shells undergoing deflation at a rate of 20 Pa/s for various hold times: 6 s (n=12), 60 s (n=13), and 600 s (n=13). Data in **c** and **i** are presented as medians with boxes indicating the interquartile range (25th to 75th percentiles).

**Figure 2:**
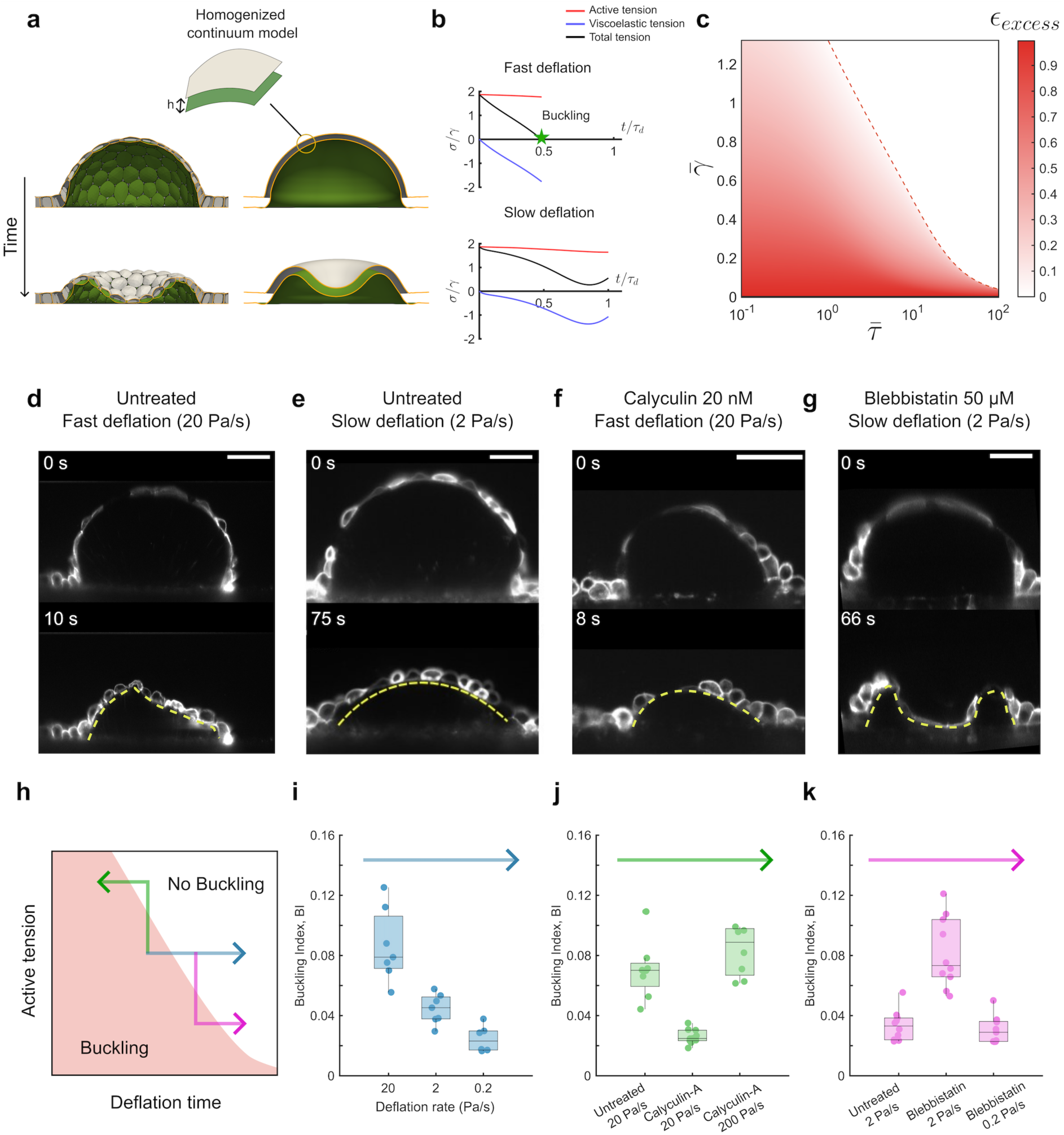
Epithelial shell buckling is determined by deflation rate and active surface tension. **a,** Illustration of the continuum model for epithelial shells, which homogenizes the 3D active gel vertex model, and where apical and basal surfaces are offsets of the mid-surface of the shell (see Supplementary Note 1). **b,** Evolution of the active, viscoelastic, and total tissue tensions *σ* normalized by the cortical active tension *γ* during fast (top) and slow (bottom) deflation relative to viscoelastic timescale. **c,** Phase diagram mapping normalized excess area ɛ_excess_ after deflation as a function of the elastocapillary parameter (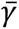 = *γ*/*μ* where *μ* is the cortical elastic shear modulus) and deflation timescale relative to viscoelastic timescale (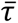 = *τ*_d_/*τ*_ve_). **d-g,** Representative images of the shell’s mid-section undergoing deflation towards a constant pressure (−50 Pa), with varying deflation rates (20 and 0.2 Pa/s) and experimental conditions (Untreated, 50µM Blebbistatin, and 20nM Calyculin). Panels **d** and **e** illustrate the deflation process for untreated shells with different deflation rates of 20 Pa/s **(d)** and 2 Pa/s **(e)**. Panel **f** demonstrates deflation under the treatment with 20nM Calyculin at a fast rate (20 Pa/s), while Panel **g** shows the deflation process under treatment with 50µM Blebbistatin at a slow rate (2 Pa/s). Yellow dashed line highlights the wrinkled or smooth curvature of the tissue section. Scale bars in **d-g**, 20 µm. **h,** Schematic illustrating the phase space explored in the experiments. **i,** Buckling index for different deflation rates (20, 2, and 0.2 Pa/s) after a hold time of 600 seconds in the untreated condition (blue path in **h**). **j,** Buckling index of the untreated condition at fast (20 Pa/s) deflation rate compared with conditions treated with Calyculin at both fast (20 Pa/s) and faster (200 Pa/s) deflation rates (green path in **h**). **k,** Buckling index of the untreated condition at slow (2 Pa/s) deflation rate compared with conditions treated with Blebbistatin at slow (2 Pa/s) and slower (0.2 Pa/s) deflation rates (pink path in **h**). Data are presented as median and box is drawn between 25 and 75 percentiles of n = 7 (untreated 20 Pa/s **(i)**), 6 (untreated 2 Pa/s **(i)**), 6 (untreated 0.2 Pa/s **(i)**) 8 (untreated 20 Pa/s **(j)**), 10 (Calyculin 20 Pa/s **(j)**), 8 (Calyculin 200 Pa/s **(j)**), 10 (untreated 2 Pa/s **(k)**), 10 (Blebbistatin 2 Pa/s **(k)**), 9 (Blebbistatin 0.2 Pa/s **(k)**).

According to this physical picture, a cell monolayer held inflated for a short time *τ*_h_relative to *τ*_ve_ should behave like an elastic balloon, with deflation merely relaxing the positive elastic tensions accumulated during inflation. By contrast, if the system was held stretched for longer times than *τ*_ve_, the ensemble of cortical surfaces should have enough time to remodel and conform to the inflated configuration, so that the reference state of the tissue is the curved epithelial shell and not the initially flat epithelium. In these conditions, fast deflation should result in compressive elastic tensions and possibly buckling, as in depressurized inert shells^5,6^.

To further examine this idea, we developed a computational framework capturing the hierarchical organization of the epithelium and bridging cytoskeletal dynamics and tissue mechanics. Briefly, the geometry of individual cells is described by triangulated surfaces enclosing cells of constant volume. Each apical, basal and lateral cell surface is modeled as an active gel capturing contractile active tension *γ*_a_ and the short-term non-linear elasticity of the network, characterized by a shear modulus integrated over the cortical thickness *μ*, and the dissipation of elastic tensions over a viscoelastic time *τ*_ve_ (Fig. 1d,e, Supp. Note 1)^26^. The effect of the ensemble of active gel surfaces enclosing fixed cellular volumes determines the emergent time-dependent mechanics of in silico epithelial shells, in particular the surface tension of the tissue, split into active and viscoelastic components, *σ*_t_ = *σ*_a_ + *σ*_ve_. This 3D active gel vertex model confirmed our rationale by predicting that epithelial shells buckle upon sudden deflation only if the hold time *τ*_h_ is longer than *τ*_ve_ (Fig. 1g, Movie 1). The wrinkles in deflated, buckled shells require excess area relative to the footprint area. To quantify it, we defined the dimensionless parameter ɛ_excess_ = *A*⁄(*πa*^2^) − 1, where *A* is the area of the monolayer and *a* is the footprint radius. ɛ_excess_ is 0 when the planar state is recovered upon deflation and ∼1 when a hemispherical purely elastic shell is deflated. Our simulations show ɛ_excess_ is 0 when the hold time is below a buckling threshold close to *τ*_ve_, and then increases and plateaus below 1 for hold times above ≈ 50 *τ*_ve_ (Extended Fig. 3).

**Figure 3:**
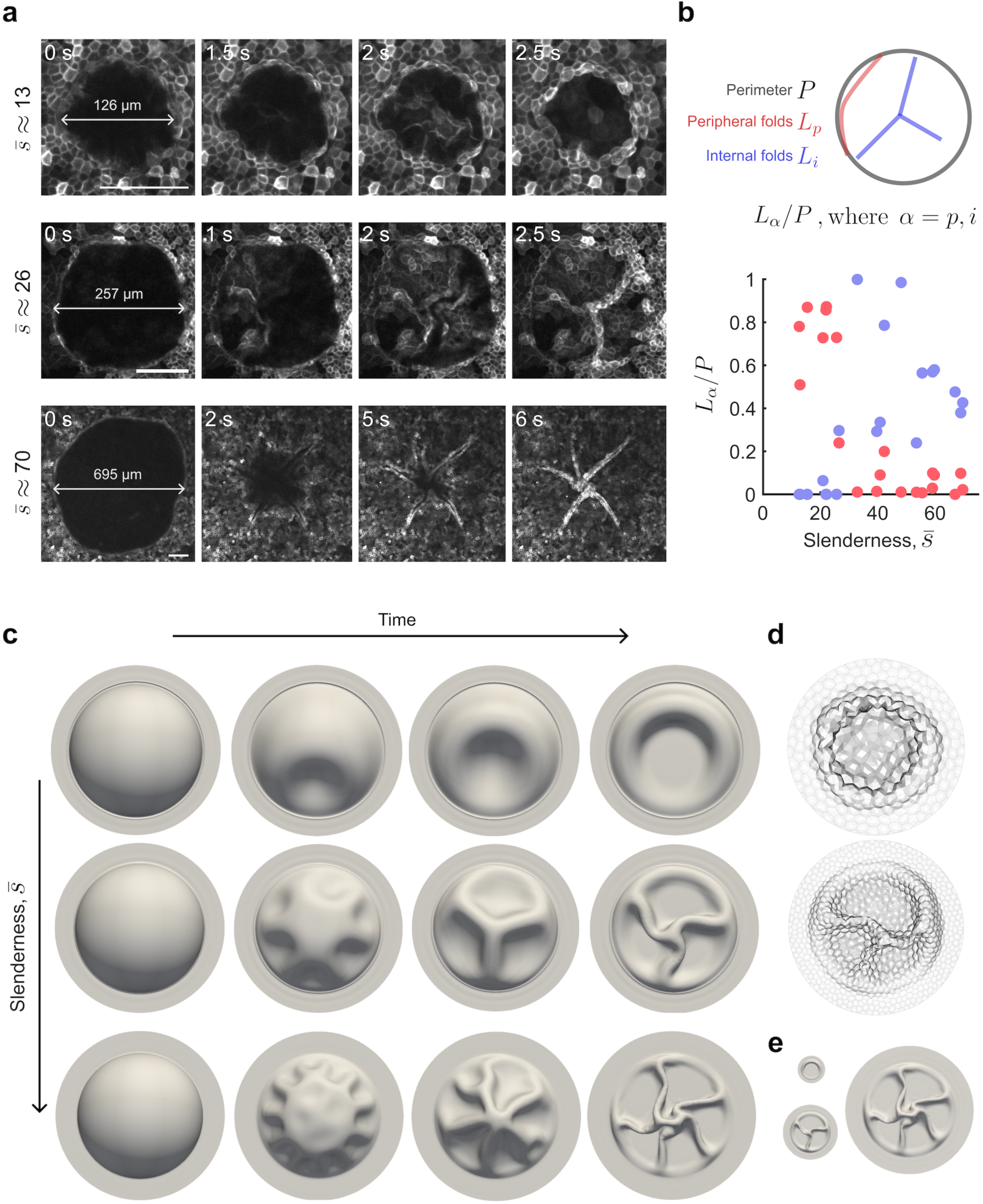
Wrinkle patterns and symmetry-breaking of epithelial shells. **a,** Confocal images of representative shells of different slenderness (*s̄* = 2*a*/ℎ_0_) undergoing buckling: *s̄* ≈ 13 (top), *s̄* ≈ 26 (middle), *s̄* ≈ 70 (bottom), (Scale bars, 100 µm). **b,** Illustration of the quantification of peripheral (red) and internal (blue) wrinkle lengths, normalized by the footprint perimeter (top). *L*_*α*_/*P* as a function of slenderness for internal and peripheral wrinkles (bottom) (n = 23). **c**, Continuum simulations of fast deflation (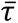 = 0.6, 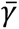 = 0.5) of epithelial shells following a long hold for the same three slenderness values as in (**a**) and for a fixed viscous drag so that *a*/*ℓ* = 0.75 (top), *a*/*ℓ* = 1.5 (center) and *a*/*ℓ* = 4 (bottom). The transition from peripheral to internal wrinkles takes place for *a*/*P* ≈ 1. **d**, Final snapshot of the same simulations using the 3D active gel vertex model. The simulation corresponding to *s̄* ≈ 70 was too expensive to run within a reasonable time. **e**, Final snapshots of (c) represented at the same scale.

To test these predictions experimentally, we formed epithelial shells on circular footprints of 80 micron diameter by applying 200 Pa for varying hold times (6, 60, and 600 s), selected based on the computational results estimating *τ*_ve_ ≈10 s^22,25,27^. We then reduced pressure down to −50 Pa at a fixed deflation rate of 20 Pa/s. To examine the deflation dynamics with high temporal resolution, we tracked the shell middle cross-section (Fig. 1h and Extended Fig. 4a) and defined a buckling index (*BI*) quantifying the presence of morphological irregularities indicative of buckling events (methods). *BI* is a proxy for ɛ_excess_, which is not experimentally accessible at rapid deflation rates. In agreement with model predictions, *BI* progressively increased as hold times were made longer, with nearly all shells exhibiting prominent buckling for the longest hold time *τ*_h_ (Fig. 1h,i and Movie 2, 3). Together, our theoretical and experimental results show that conditioning epithelial sheets to long periods of inflation is essential to dissipate viscoelastic tensions and hence generate curved epithelial shells that buckle when deflated. We hence focus only on epithelial shells held inflated for 600 s in the remainder of the paper.

**Figure 4:**
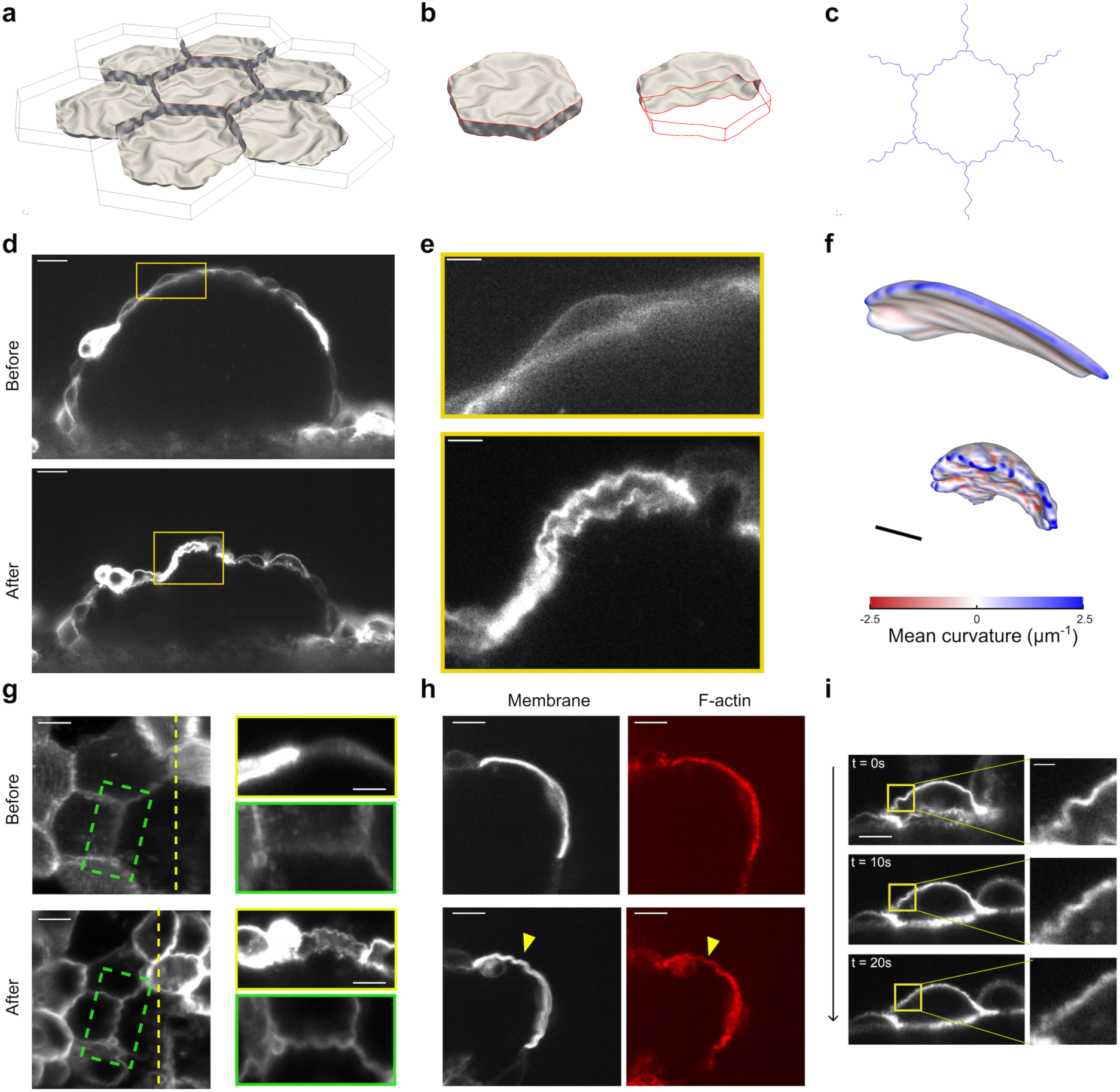
Multiscale buckling of epithelial shells. **a-c,** Simulations of equibiaxial compression of an epithelial patch in an active gel vertex model explicitly accounting for cytosolic viscous hydrodynamics and cortical bending rigidity. The dashed polygons indicate the epithelial patch before compression (see also Movie 9). **(a)**, View of 7 cells where some surfaces have been removed for visualization, **(b)** cell in the center, **(c)** cross section of lateral surfaces. **d,** Shell’s mid section before (top) and after (bottom) buckling, obtained with light sheet microscopy (Scale bar, 20 µm). **e,** Zoomed inset of the region marked with a yellow box in (d), highlighting subcellar wrinkles (representative of 25 shells, Scale bar, 5 µm). **f,** The same cell is rendered in 3D as a closed surface, color coded for mean curvature (see also Movie 10). **g,** Left, maximum intensity projection of the epithelium before (top) and after (bottom) the buckling event. Right, yellow rectangles: lateral view along the dashed yellow line marked on the left panels, revealing both apical and basal surface buckling. Right, green rectangles: zoomed view of a junctional buckling event corresponding to the region marked in green in the left panel (representative of 8 shells, Scale bar, 5 µm). **h,** Cross-section of a cell undergoing subcellular buckling, showing the membrane (left, CIBN-GFP-CAAX) and F-actin (right, SPY-actin) (n = 10; Scale bar, 5 µm) **i,** Timelapse of a buckled cell immediately after deflation, with an inset highlighting the relaxation of the cortical fold (n = 6, Scale bar, 5 µm).

### Parameters controlling buckling of epithelial shells

To identify the mechanical parameters controlling the emergence of buckling in our system, we resorted to continuum theories for epithelial shells enabling a more systematic parameter exploration^28–30^. For this purpose, we developed a nonlinear continuum model that homogenizes the active gel vertex model of the tissue by effectively accounting for the ensemble of the cortical active viscoelastic surfaces, and where apical and basal surfaces are offsets of the mid-surface of the shell of thickness ℎ (Fig. 2a, Supp. Note 1)^31^. In addition to enabling more efficient simulations while accurately conforming to the 3D active gel vertex model (Fig. 2a, Extended Fig. 5, Movie 1), the continuum model identifies a minimal set of dimensionless parameters controlling the system behavior: the slenderness of the shell defined as *s̄* = 2*a*/ℎ_0_, where ℎ_0_ ≈ 10 μm is the cell thickness before inflation and *a* the footprint radius; the ratio between the deflation time and the viscoelastic time, 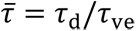; an elastocapillary parameter^32^ given by the ratio between the average apicobasal active tension 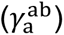 and shear elastic modulus of the cortex surface, 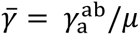; and a dimensionless prestress parameter combining the aspect ratio of cells in the planar state and the ratio of lateral to apicobasal active tensions, 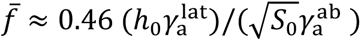, where *S*_0_ is the typical apical area of cells before inflation.

**Figure 5:**
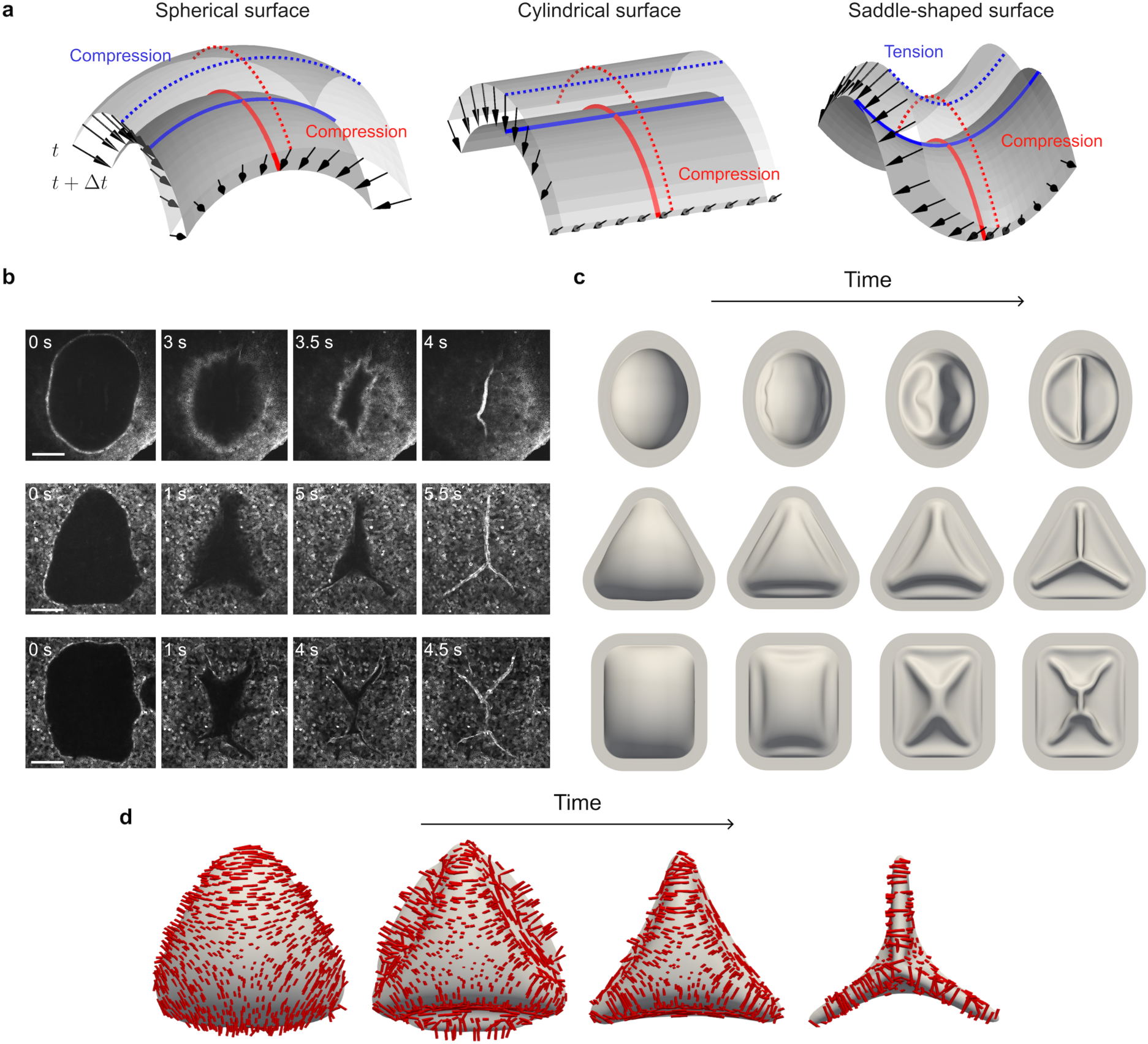
Geometric control of wrinkle patterns. **a**, Illustration of how anisotropic curvature induces anisotropic strain rate for three surfaces undergoing normal motion representative of deflation, with spherical surfaces being compressed in all directions, cylindrical surfaces being anisotropically compressed, and saddle-shaped surfaces being compressed or stretched depending on the direction. **b**, Confocal images of shells undergoing buckling, showing different footprint shapes: ellipse (top), triangle (middle), and rectangle (bottom) (Scale bars, 100 µm). **c**, Continuum simulations corresponding to (c) (*s̄* ≈ 40, 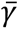 = 0.5, 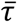 = 0.6, *a*/*ℓ* ≈ 4). The side views in Movie 12 show that peripheral folds are very shallow compared to internal folds. **d**, Curvature anisotropy in the top part of the epithelial shell with triangular footprint during deflation. The red segments show the direction of maximum curvature, and their length is proportional to the difference between the principal curvatures.

According to the model, *σ*_a_ slightly decreases during deflation but remains positive (Fig. 2b). By contrast, *σ*_ve_, which is initially zero after a long hold time, becomes negative as the tissue becomes laterally compressed and relaxes at a rate proportional to *σ*_ve_/τ_ve_. Hence, the total tension in the tissue *σ*_t_ may become negative, leading to buckling. However, whether this critical point is attained before complete deflation depends on dimensionless parameters 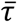 and 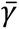; both larger deflation times, enabling viscoelastic tension relaxation, and higher active tensions, compensating negative viscoelastic tensions, tend to suppress buckling. This behavior is mapped by a phase diagram identifying the buckling domain in the parameter plane 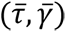, characterized by the condition *σ*_t_ < 0 being met prior to full deflation, along with the normalized excess area in the tissue after full deflation ɛ_excess_ (Fig 2c, Supp. Note 2 and Extended Fig. 6).

To test this conceptual framework for epithelial shell buckling, we designed experiments to explore different paths within the phase diagram (highlighted in different colors in Fig. 2h). First, we conducted experiments at different deflation rates, resulting in deflation times of about 10, 100 and 1000 s (blue path in Fig. 2h). In agreement with the theory, we found that buckling could be largely reduced by increasing the deflation time (Fig 2e,i, Extended Fig. 4b, Movie 3). Besides deflation time, our theory predicts that buckling can also be reduced by increasing surface tension (green path in Fig. 2h). To test this prediction, we focused on fast-deflation conditions and treated monolayers with Calyculin-A, known to enhance myosin II activity by inhibiting myosin phosphatase. As predicted, we found a significant reduction of *BI* (Fig 2f,j, Extended Fig. 4c, Movie 4). Moreover, we confirmed that under these conditions of high surface tension, buckling could be restored by further increasing deflation rate (Fig. 2j). Conversely, we experimentally inhibited contractility by treating our monolayers with Blebbistatin, which reduces active tension by blocking myosin ATPase activity (violet path in Fig. 2h), observing increased buckling at intermediate deflation rates, which as predicted could be reduced by slowing down deflation (Fig. 2g,k, Extended Fig. 4d, Movie 4). In summary, our experiments confirmed a physical picture according to which the negative elastic stresses in compressed epithelial shells can be buffered by either viscoelastic relaxation or by buckling, with this competition being controlled by active contractility and loading rate (Fig. 2h). Comparison between experiment and theory allowed us to estimate the order of magnitude of the key parameters in the model (Supp. Note 3). Of note, our estimate for the elastocapillary parameter, 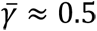,^27,33^ is orders of magnitude larger than that of various inert thin films interacting with fluids^32^, suggesting that epithelial shells probe a rather unusual buckling regime characterized by large surface tension, allowing the compressed cellular sheet to significantly reduce its area before buckling.

### Symmetry-breaking of wrinkle patterns

Wrinkling patterns in inert shells are known to strongly depend on slenderness^4,6^. Accordingly, we performed rapid deflation experiments on epithelial shells with footprint diameters ranging from 100 μm to 1 mm, corresponding to a slenderness parameter *s̄* = 2*a*/ℎ_0_ between 10 and 100. Using confocal microscopy, we examined the geometry of the wrinkle patterns at the base of the deflated shell (Fig. 3a, Movie 5). We found that epithelial shells of low slenderness systematically developed peripheral cell accumulation, whereas more slender shells broke axisymmetry and formed networks of relatively straight internal wrinkles, often arranged radially. To quantify this transition, we measured the ratio *L*_*α*_/*P*, where *L*_*α*_ is the wrinkle length (with the subscript *α* denoting wrinkle localization: *i* for internal and *p* for peripheral), and *P* is the footprint perimeter (Fig. 3b). This ratio indicates the extent of wrinkling relative to the dome footprint size and whether the wrinkles are predominantly internal or peripheral. The transition between peripheral and internal wrinkles occurred around a critical slenderness parameter *s̄*_crit_ ≈ 30, that is for diameters of around 300 μm (Fig. 3b). The wrinkles remained imprinted into the cell monolayer for tens of hours, a much longer time-scale than that of the sub-cellular processes controlling their formation (Extended Fig. 7), likely due to the formation of adhesive links between the cell monolayer and the substrate.

To understand the transition between peripheral and internal wrinkles, we turned to our theoretical model. Previous work on deflated spherical elastic shells has shown that at small deflation or for thick shells, the buckling mode consists of an axisymmetric dimple where curvature is inverted, the Pogorelov buckling mode (Fig. 2a), whereas thinner shells under larger deflation break axisymmetry to develop a wrinkled polygonal dimple^18,34,35^. In other situations, the geometric frustration of a thin spherical shell confined to a plane results in low-symmetry patterns of internal wrinkles^36,37^. Our continuum simulations reproduced the smooth and polygonal Pogorelov dimples depending on slenderness and degree of deflation, but these modes always resulted in peripheral area accumulation and failed to capture the internal wrinkles observed experimentally (Movie 6). We then hypothesized that these internal wrinkles could arise from the drag associated to displacing the viscous medium surrounding the epithelial shells. Indeed, it is widely appreciated that when a plate supported by a soft elastic material is compressed, the competition between elastic deformation in the plate and the substrate selects the surface wrinkle wavelength^11,14,38^. Analogously, the competition between elastic energy release in the plate and dissipation in a surrounding viscous^38–41^ or viscoelastic^42,43^ medium can also result in pattern selection at the onset of wrinkle formation, often followed by pattern coarsening. Following this rationale, we used scaling arguments to estimate the dominant wavelength at the onset of the buckling pattern as

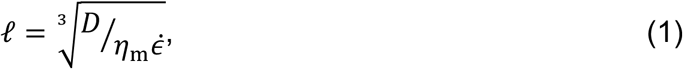

where *D* is the bending rigidity of the shell, *η*_m_ is the viscosity of the medium and 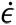 is the strain rate (Supp. Note 4). According to this scaling, if the system size is larger than *P*, then a buckling pattern with internal structure should develop, possibly leading to internal wrinkles that break symmetry as a result of geometric frustration. The critical slenderness for the transition from peripheral to internal wrinkles follows as 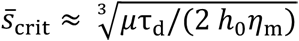. Reasonable parameters lead to *s̄*_crit_ ≈ 37, in the same order of magnitude as our experimental results (Fig. 3b, Supp. Note 4), supporting the idea that viscous drag may underlie symmetry breaking.

To test this idea further, we included a viscous drag of the embedding medium in our simulations. We first fixed the size *a* of the shell footprint and varied the magnitude of the viscous drag, and hence of the dominant wavelength *ℓ*. In agreement with the theoretical argument, when the system size relative to the dominant wavelength (*a*/*ℓ*) is small, a single dimple develops and deepens during post-buckling, leading to excess area accumulation at a peripheral wrinkle. Conversely, for large *a*/*ℓ*, the system first develops a pattern of dimples whose size closely follows *ℓ*, which then coarsens upon further deflation leading to a network of internal wrinkles whose complexity increases with *a*/*ℓ*. The transition takes place when *a*/*ℓ* ≈ 1 (Extended Fig. 8 and Movie 7). To recapitulate the experimentally observed transition from peripheral to internal wrinkles at a critical slenderness, we varied the system size and fixed the drag coefficient, resulting in *a*/*ℓ* ≈ 0.75; 1.5; 4. Both the 3D active gel vertex model and the continuum model closely matched the experimental post-buckling patterns, and further provided a mechanistic pathway from the pattern of dimples at buckling onset to the final wrinkles (Fig. 3c-e, Extended Fig. 9 and Movie 8). Taken together, these results support that viscous confinement by the surrounding medium controls the peripheral-to-internal transition as system size increases.

### Hierarchical buckling at multiple scales

We then reasoned that the arguments leading to Eq. (1) for the dominant wavelength of a dynamically compressed epithelial shell are generally applicable to other active surfaces in the system, and particularly to the actomyosin cortex. Indeed, the cortex is an active viscoelastic shell surrounded by a highly viscous cytosol. Using estimates of cortical thickness^16^ (100 nm) and cytosol viscosity^44^ (0.6 Pa·s), our scaling arguments predict the emergence micron-sized cortical wrinkles under fast deflation (Supp. Note 4). We confirmed this estimation by performing fast compression simulations of epithelial monolayers with a detailed active gel vertex model accounting for the bending rigidity of cortical surfaces and for cytosolic viscous hydrodynamics (Fig. 4a-c, Supp. Note 5 and Movie 9).

To test this prediction experimentally, we used fast light sheet microscope to obtain snapshots of the shells with sufficient time and space resolution to detect cortical wrinkles. As predicted, we found that tissue-scale buckling coexisted with micron-scale wrinkling of apical, basal and lateral surfaces (Fig. 4d-g, Movie 10). Simultaneous imaging of the membrane and the cortex showed that both structures co-localized throughout the buckling process, ruling out that wrinkles could originate from membrane detachment, Fig. 4h. As expected, subcellular wrinkles were short-lived, dissipating within ∼20 s (Fig. 4i). Taken together, our results portray epithelial tissues as active hierarchical materials able to relax compressive stresses by wrinkling at multiple spatiotemporal scales.

### Engineering epithelial wrinkles by geometry

The wrinkle patterns obtained so far result from symmetry breaking of axisymmetric shells and thus, aside estimations of their characteristic lengths, the precise location of the final wrinkles will depend on unpredictable fluctuations in the system. We next sought to guide the wrinkle patterns deterministically. We reasoned that this could be achieved by introducing compression heterogeneity to define nucleation sites, and anisotropy to orient wrinkles perpendicular to the direction of maximal compression.^45^. In our system, we note that the in-plane strain rate in a curved shell due to shape changes can be expressed as −*v*_*n*_***k***, where *v*_*n*_ is the normal velocity of the deflating shell and ***k*** is the extrinsic curvature tensor of the shell surface^46^ (Fig. 5a). Therefore, curvature anisotropy implies compression anisotropy.

To generate anisotropic curvature in MOLI, we formed large epithelial shells with footprints shaped as ellipses, triangles and rectangles (Fig. 5c). Upon rapid deflation, we observed a highly reproducible arrangement of internal wrinkles that depended on footprint geometry. Specifically, elliptical footprints led to wrinkles along the long axis, while polygonal footprints gave rise to internal wrinkles emanating from the corners. In all cases, wrinkles conformed to the medial axis of the footprint, defined as the set of points having more than one closest point to the boundary (Fig. 5c and Movie 11). The medial axis has been shown to organize winkles in positively curved inert thin sheets confined to a plane^36,37^. However, in contrast to our system, those wrinkles form perpendicular to the medial axis^36,37^, which we attribute to stark differences in *s̄*, 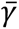, curvature and boundary conditions. In our system, simulations show that the curvature anisotropy along the footprint edges leads to peripheral inward wrinkles, which progressively deepen and define lines that approximate the medial axis, along which curvature anisotropy causes transversal compression and wrinkling (Fig. 5b,d, Movie 12). Despite the fact that here *a*/*ℓ* is significantly greater than 1, buckling does not lead the emergence of a pattern of dimples as in spherical shells, and instead it is directed by geometry. In summary, the control of pressure and footprint shape provided by MOLI determine the magnitude, rate and anisotropy of compression, enabling the deterministic programming of epithelial wrinkles.

### Discussion and outlook

Our study delineates the potential of epithelial buckling as a morphogenetic mechanism and establishes its mechanical rules. The combination of our new experimental approach with computational models bridging cytoskeletal and tissue dynamics allowed us to systematically map a previously unexplored regime of buckling dynamics across a broad range of time scales, length scales, geometries, and mechanical properties of the epithelium. Being devoid of extracellular matrix, our epithelial shells allow us to focus on the intrinsic mechanics of the cell monolayer. Beyond the widely invoked scenario of buckling in a laterally compressed monolayer^14,47–50^, our analysis establishes the concept of active viscoelastic buckling as a morphogenetic mechanism, which may play a role in morphogenetic processes with time scales comparable those of cortical relaxation, such as those involving rapid contractions, lumen bursting or tissue fracture^51–53^. This mechanism offers a new regulatory layer for buckling morphogenesis where folds and wrinkles can be controlled, not only through the rate of compression, but also through pre-existing strain conditioning or surface tension in the tissue.

Besides establishing the mechanisms that govern the onset buckling, our work also characterizes the post-buckling patterning of wrinkles. We have shown that wrinkling organization arises from an interplay between mechanical confinement by the surrounding medium, which controls the wrinkle density, and shell geometry, which induces frustration and anisotropic compression. Interestingly, our results on cortical buckling indicate that analogous rules operate at two different scales in the hierarchical organization of cell monolayers. With this understanding, we were able to harness the active viscoelasticity of the tissue and footprint geometry to rationally design long-lasting wrinkling patterns. We showed that wrinkles follow the medial axis of polygonal footprints, providing a simple geometric rule for the generation of stable wrinkles in cell monolayers. This approach can be used to build functional structures for applications in synthetic morphogenesis, for example to engineer perfusable networks for vasculogenesis^54^.

## Methods

### Replica molding of Polydimethylsiloxane gels

Polydimethylsiloxane (PDMS) (Sylgard PDMS kit, Dow Corning) was utilized to fabricate microfluidic devices by mixing the curing agent and elastomer in a 1:9 weight ratio. This mixture was centrifuged for 2 min at 900 rpm to eliminate air bubbles. The unpolymerized PDMS was subsequently poured into a mold to obtain the desired shape.

### Fabrication of microfluidic devices

We employed two distinct types of devices. One was designed for compatibility with an inverted microscope (TiE, Nikon) equipped with a confocal unit (CSU-W1, Yokogawa), and the other was intended for use with a dual-view upright light sheet microscope (QuVi SPIM, Luxendo Light-sheet microscopes, Bruker).

The device designed for inverted confocal microscopy consists of four distinct components. The first component is a PDMS top block, 8 mm thick, four inlets, and a channel intended for the application of hydraulic pressure (Extended Figure 1). The second component is a 200 µm thin PDMS layer with a 1.2 mm diameter hole at the center covered with a porous membrane (Polycarbonate filtration membrane, pore size: 400 nm and 3 µm, Whatman). The third component is another PDMS layer, which can be either 200 µm or 500 µm thin, and includes a channel specifically designed for cell seeding. The thickness of this layer is set according to the shell size. Finally, all these PDMS components are assembled on the fourth component, a glass-bottomed dish (35 mm, 0 coverslip thickness, Cellvis), serving as a container for the device.

The top block was fabricated through replica molding, utilizing a 3D printed mold created with vat polymerization and a digital light processing 3D printer (Solus DLP 3D Printer with SolusProto resin, Reify 3D). Following this, the mold’s surface was treated with Trichlorosilane (Trichloro(1H,1H,2H,2H-perfluorooctyl)silane, Merck) to ensure that it does not adhere to PDMS. Unpolymerized PDMS was then poured into the mold and degassed for one hour. Subsequently, the PDMS was cured at 100°C on a hot plate for 30 minutes. After curing, the PDMS was carefully removed from the mold, cut into the device shape, and punched with 1.5 mm holes in the inlets.

The thin PDMS layers were produced by pouring unpolymerized PDMS onto a 15 cm plastic dish. The thickness of these layers is determined by the volume of PDMS used: 4 ml for a 200 µm layer and 10 ml for a 500 µm layer. These dishes were then polymerized in an oven at 75°C for 12 hours. Following polymerization, the thin sheets were cut into the appropriate device parts using a Silhouette cutting machine (Silhouette Cameo 4, Silhouette America). The sheets were carefully peeled off the plastic dish and placed on a Silhouette cutting mat. The Silhouette cutting machine was used to cut device layers in PDMS sheets according to patterns specified in Silhouette software.

The assembly of the devices was facilitated by an ozone plasma cleaner (PCD-002-CE, Harrick Plasma). The glass-bottomed dishes and thin PDMS layers with cell channels were subjected to plasma treatment for 1 minute, followed by bonding the layers together by placing them in contact for 2 hours at 75°C. The top block and the middle layer were bonded using the same method. For the attachment of porous membranes to PDMS layers, the membranes were first plasma-treated for 1 minute. Subsequently, they were treated with (3-aminopropyl) triethoxysilane (APTES, Sigma-Aldrich, cat. no. A3648), diluted at 5% in MilliQ water solution at 80°C for 20 minutes. The top block, which was bonded to the middle layer, was treated with plasma treatment for 1 minute to bond the porous membrane. Finally, all these parts were bonded together once more using the same plasma bonding protocol.

The devices designed for light-sheet microscopy were constructed from a single PDMS block bonded to a glass microscope slide (76×26 mm, BPB016, RS Components). These blocks were fabricated using a 3D printed mold (Ultimaker 3 with Ultimaker PLA Printer Filament 1616, Ultimaker). The PDMS was mixed, centrifuged, degassed, and cured as previously described. After curing, the PDMS was carefully removed, cut into individual devices, and holes were punched with a 1.5 mm biopsy punch. The PDMS blocks were then bonded to glass slides by applying a thin layer of unpolymerized PDMS, which was spread onto the glass slides using a spatula. Following this, the devices were placed on a hotplate at 100°C for 30 minutes to complete the bonding. Subsequently, the porous membranes were attached to the devices using same protocol as described previously.

### Patterning protein on the devices

The devices were initially filled with 96% ethanol to remove air bubbles. Subsequently, they were filled with MilliQ water to eliminate ethanol traces. The Light-Induced Molecular Adsorption of Proteins technique (PRIMO, Alveole Lab) was employed to pattern adhesion-promoting protein. For this procedure, the devices were incubated with 1% Poly-L-lysine (PLL) (P2636, Merck) diluted in type 1 water for 1 hour, followed by 50mg/ml PEG-SVA (Laysan Bio) in 8.24pH 10mM HEPES (1M, Merck) for 30 minutes, and then rinsed with HEPES. Prior to using PRIMO, the devices were filled with a photoinitiator, P-benzoylbenzyl trimethylammonium chloride (Merck). The desired protein pattern was loaded into the PRIMO software (Alveole Lab, Leonardo), which projects UV (375 nm) light over the porous membrane. The UV light pattern (dose 1500 mJ/mm^2^) cuts the PEG chains, enabling the attachment of adhesion-promoting proteins. After the PRIMO process, the samples were rinsed with phosphate-buffered saline (PBS, Merck). The samples were then filled with fibronectin (F0895, Sigma-Aldrich) and fibrinogen (100 µg/ml Fibronectin in 2% Far-red (647 nm) fibrinogen solution in 1X PBS) for 5 minutes. The samples were rinsed again with 1X PBS. Fibrinogen with a far-red signal was used to image the coated protein pattern, enabling the tracking of the epithelial shell positions. The samples were stored at 4°C for two to three days before cell seeding.

As the PRIMO system is mounted on an inverted confocal microscope, devices intended for light sheet microscope needed special adaptations. A PDMS spacer and glass coverslip are utilized to form a temporary covered channel over the porous membrane, allowing the use of an inverted microscope by flipping the device upside down. This spacer accommodates the photoinitiator during the PRIMO process. The spacer is fabricated from 2 × 2 cm squares of a 400 µm thick PDMS layer, with a keyhole shape cut from the side. Each PDMS piece is then adhered to a glass coverslip (18 mm, 1# Cover glasses circular, 0111580, Marienfeld) using the surface tension of the liquid.

### Cell culture in the device

To image cell shape and tissue structure, experiments were conducted using MDCK strain II line cells expressing CIBN-GFP-CAAX, which labels the plasma membrane (kindly provided by Guillaume Charras). These cells were cultured in Dulbecco’s Modified Eagle Medium (DMEM, Gibco Thermofisher) supplemented with 10% v/v fetal bovine serum (FBS, Gibco, Thermofisher), L-glutamine (Thermofisher), 100 µg/ml streptomycin, and penicillin. The cells were incubated at 37°C under a 5% CO_2_ condition. Prior to seeding cells in the device, it was filled with cell culture medium. The cells were trypsinized and diluted to a concentration of 25 ─ 30×10^6^ cells/ml. For inverted microscope devices, the cell channel of the device was filled with 30 µl of cell solution and incubated to allow cell adhesion. For upright microscope devices, 30 µl of cell solution were added using the spacer. After one hour of incubation, the devices were rinsed with media to remove unattached cells. The devices were then incubated for 24 hours in the incubator to allow the growth of a monolayer before the experiment.

### Labelling actin with SPY-actin

To observe the dynamics of the cortex, we utilized SPY555-actin (Spirochrome), a bright dye specifically optimized for the rapid labeling of F-actin in live cells with minimal background. To prepare the 1000x solution, we added 50µl of anhydrous DMSO to the stock SPY555-actin. Subsequently, we added 1µl of the 1000x solution to 999µl of cell culture medium. The resulting solution was then introduced into the microfluidic chips and incubated for two hours prior to imaging.

### Alteration of active tension

To increase active tension, 20nM Calyculin A was used to enhance Myosin II activity by inhibiting Myosin phosphatase (MLCP). Conversely, 50µM Para-Nitroblebbistatin was used to reduce active tension by blocking myosin ATPase activity. All pharmacological treatments were done with 20 min incubation before application of pressure.

### Application and measurement of the pressure

The tissue is cultured on porous membranes and inflated into epithelial shells by applying pressure beneath the membrane through a channel. This channel features one inlet and one outlet for bubble removal. The inlet is connected to a 35 ml reservoir of cell culture medium (in a 50 ml falcon tube) via PTFE tubing (1/16-inch OD for Microfluidics, Darwin microfluidics), and the outlet is connected to a shutoff valve (Microfluidic Sample Injection / Shut-off Valve, Darwin microfluidics). All tubes are connected to the chip with a steel insert (Stainless steel 90° Bent PDMS Couplers, Darwin microfluidics). Once bubbles are removed by passing cell culture medium, the valve is closed to allow pressure application on the basal side of the cells. Pressure is measured by comparing the vertical position of the sample and the liquid-air interface in the reservoir. To induce precise pressure changes, a motorized linear translation stage (Zaber) is used to lift the fluid reservoir. The Zaber software facilitated both the measurement and regulation of the pressure protocol, ensuring accurate and controlled pressure conditions throughout the experiments.

### Confocal Microscopy

For 3D timelapse imaging of shells over larger time intervals (> 1 min), an inverted Nikon microscope equipped with a spinning disk confocal unit (CSU-W1, Yokogawa) was utilized, along with Nikon 40× 0.75 NA, 20× 0.75 NA, and 10× 0.45 NA air lenses. For shorter time intervals (< 10 s), a Zeiss LSM880 inverted confocal microscope was employed in laser scanning mode. Fast imaging was facilitated by imaging a single line of pixels passing through the center of the shell footprint.

### Light-sheet microscopy

Light sheet imaging was conducted using a dual-illumination inverted Selective Plane Illumination Microscope (diSPIM) (QuVi SPIM, Luxendo, Brucker), equipped with Nikon 40x immersion lenses (Nikon CFI Apo 40× W 0.8 NA NIR water immersion objective). For fast buckling experiments, single objective illumination and detection were used at a frame rate of 0.5 frames per second.

### Quantification of the shell areal strain and tension

ImageJ was used to manually section the shell along the XZ plane (the XY plane is the plane parallel to the flat monolayer.) This section was used to calculate the shell’s height *h*, radius of curvature *R*, and base radius *a*. Strain *ɛ* and tension *σ* were calculated using the applied pressure values *ΔP* with the following equations:

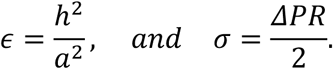

The raw data was extracted using ImageJ 1.53t and then processed in MATLAB 2023a (MathWorks) to compute and plot the strain and tension.

To capture fast dynamics, a single XZ plane passing through a diameter of the shell footprint was imaged over time. This time-lapse was used to create kymographs representing changes in height over time. These kymographs were saved as image files and later analyzed using MATLAB. A custom-built MATLAB code was used to digitize the kymographs, where the maximum intensity along each time point was considered as the current shell height position. The shell height was calculated as the difference between the current shell height position and the initial (non-inflated) position. The radius of curvature was calculated using the relation:

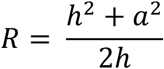

### Analysis of the buckling event

To quantify the extent of buckling, the shell outline was manually segmented at the basal side of the tissue for different time points during the deflation. The curve obtained was used to calculate the area under the curve (*A_curve_*) and its convex hull area (*A_hull_*). A buckling index (*BI*) was defined as the maximum difference over time between *A_hull_* and *A_curve_*, normalized by the maximum of *A_curve_* (*A_0_*).

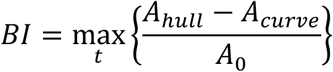

In non-buckling cases, the difference between *A_hull_* and *A_curve_* is small, as the shell maintains uniform curvature, resulting in a low *BI*. Conversely, in buckling cases, *A_hull_* increases compared to *A_curve_*, leading to a higher *BI*.

### Qualitative analysis of the buckling event

In light sheet microscopy images, the buckling status of shells was determined by manually inspecting each frame during the deflation process. Shells were marked as ‘not buckling’ if they maintained a smooth circular geometry in the XZ plane throughout the deflation. Conversely, shells exhibiting a visual discontinuity in curvature or a kink were classified as ‘buckling’. The ‘buckling’ shells were analyzed by examining stacks using the ImageJ plugin 3Dscript and the Python library Napari (version 0.4.19) to identify subcellular wrinkles. The 3D reconstruction of cells was achieved by first denoising 3D images with Noise2void (https://github.com/juglab/n2v), followed by cell segmentation using a custom Python script. This script incorporated 3D watershed segmentation along with manual corrections. Finally, the segmentation mask was imported into Blender 4.0 (Blender foundation) for 3D rendering using the ‘tif loader’ add-on (https://github.com/oanegros/tif2blender).

### Quantitative analysis of the buckling pattern

To observe the buckling pattern, XY confocal images were obtained of the base of the deflating shell, near the apical surface of the monolayer, to ensure the visibility of the buckled epithelial structure. Radial and peripheral wrinkles were manually marked using a custom MATLAB script. The complexity of the pattern was quantified as the length of peripheral folds (*L_p_*) and of internal folds (*L_i_*) normalized by the perimeter (*P*) of the shell base.

## Acknowledgments

We thank all the members of our groups for their discussions and support, particularly, Özge Özguç, Ariadna Marín-Llauradó, Isabela Fortunato, Raimon Sunyer, and Alejandro Torres-Sánchez. We thank Anghara Menéndez Montes, Susana Usieto, Mònica Purciolas, and Microfabrication facility of IBEC for technical assistance. This paper was funded by the Generalitat de Catalunya (AGAUR SGR-2017-01602 to X.T., AGAUR 2021-SGR-01049 to M.A., the CERCA Programme, and “ICREA Academia” awards to P.R-C. and M.A.); the Spanish Ministry for Science and Innovation MICCINN/FEDER (PID2021-128635NB-I00, MCIN/AEI/ 10.13039/501100011033 and “ERDF-EU A way of making Europe” to X.T., PID2022-142178NB-I00 to M.A., PID2019-110298GB-I00 to P.R-C.); European Research Council (CoG-681434 to M.A., 101097753 to P.R-C. and Adv-883739 to X.T.); La Caixa Foundation (LCF/PR/HR20/52400004 to P.R-C. and X.T.); IBEC is recipient of a Severo Ochoa Award of Excellence from the MINECO. T.G. is funded by the Deutsche Forschungsgemein-schaft (DFG, German Research Foundation) – 445510097, and T.W. is funded by the Spanish Ministry of Science and Innovation (PRE2020-092774).

## Author contributions

N.C., M.A., and X.T. conceived the project. N.C. performed experiments, except for light sheet microscopy, which was performed by T.W.. N.C., analyzed data from confocal microscopy and line scan microscopy. T.W. analyzed data from light sheet microscopy. P.P.V. performed the 3D reconstruction of light sheet microscopy images. N.C., T.G., and T.W. developed protocols for the fabrication of microfluidic devices. A.O., P.B., G.V., and M.A. developed the computational framework and conducted numerical investigations. Specifically, A.O. and M.A. developed the active gel vertex model; G.V., A.O. and M.A. extended it for the cortical wrinkling simulations; and P.B. and M.A. developed the homogenized bilayer model. P.R-C. contributed technical expertise and materials. N.C., M.A. and X.T. wrote the manuscript. All authors revised the completed manuscript.

## Code availability

Analysis procedures and code implementing the model are available from the corresponding authors on reasonable request.

## Data availability

The data that support the findings of this study are available from the corresponding authors on reasonable request. Extended Data is available for this paper. Correspondence and requests for materials should be addressed to M.A. or X.T.

## References

1. Iijima, S., Brabec, C., Maiti, A. & Bernholc, J. Structural flexibility of carbon nanotubes. J. Chem. Phys. 104, 2089–2092 (1996).

2. Lidmar, J., Mirny, L. & Nelson, D. R. Virus shapes and buckling transitions in spherical shells. Phys. Rev. E 68, 051910 (2003).

3. Ribe, N. M., Stutzmann, E., Ren, Y. & van der Hilst, R. Buckling instabilities of subducted lithosphere beneath the transition zone. Earth Planet. Sci. Lett. 254, 173–179 (2007).

4. Audoly, B. & Pomeau, Y. Elasticity and Geometry: From Hair Curls to the Nonlinear Response of Shells. (Oxford University Press, Oxford, New York, 2010).

5. Bushnell, D. Buckling of Shells-Pitfall for Designers. AIAA J. 19, 1183–1226 (1981).

6. Stoop, N., Lagrange, R., Terwagne, D., Reis, P. M. & Dunkel, J. Curvature-induced symmetry breaking determines elastic surface patterns. Nat. Mater. 14, 337–342 (2015).

7. Kim, J. B. et al. Wrinkles and deep folds as photonic structures in photovoltaics. Nat. Photonics 6, 327–332 (2012).

8. Terwagne, D., Brojan, M. & Reis, P. M. Smart Morphable Surfaces for Aerodynamic Drag Control. Adv. Mater. 26, 6608–6611 (2014).

9. Yang, S., Khare, K. & Lin, P.-C. Harnessing Surface Wrinkle Patterns in Soft Matter. Adv. Funct. Mater. 20, 2550–2564 (2010).

10. Tallinen, T. et al. On the growth and form of cortical convolutions. Nat. Phys. 12, 588–593 (2016).

11. Ben Amar, M. & Jia, F. Anisotropic growth shapes intestinal tissues during embryogenesis. Proc. Natl. Acad. Sci. 110, 10525–10530 (2013).

12. Liang, H. & Mahadevan, L. The shape of a long leaf. Proc. Natl. Acad. Sci. 106, 22049–22054 (2009).

13. Yan, J. et al. Mechanical instability and interfacial energy drive biofilm morphogenesis. eLife 8, e43920 (2019).

14. Nelson, C. M. On Buckling Morphogenesis. J. Biomech. Eng. 138, (2016).

15. Wyatt, T., Baum, B. & Charras, G. A question of time: tissue adaptation to mechanical forces. Curr. Opin. Cell Biol. 38, 68–73 (2016).

16. Kelkar, M., Bohec, P. & Charras, G. Mechanics of the cellular actin cortex: From signalling to shape change. Curr. Opin. Cell Biol. 66, 69–78 (2020).

17. Carlson, R. L., Sendelbeck, R. L. & Hoff, N. J. Experimental studies of the buckling of complete spherical shells. Exp. Mech. 7, 281–288 (1967).

18. Quilliet, C., Zoldesi, C., Riera, C., van Blaaderen, A. & Imhof, A. Anisotropic colloids through non-trivial buckling. Eur. Phys. J. E 27, 13–20 (2008).

19. Quilliet, C. Numerical deflation of beach balls with various Poisson’s ratios: From sphere to bowl’s shape. Eur. Phys. J. E 35, 48 (2012).

20. Latorre, E. et al. Active superelasticity in three-dimensional epithelia of controlled shape. Nature 563, 203–208 (2018).

21. Marín-Llauradó, A. et al. Mapping mechanical stress in curved epithelia of designed size and shape. Nat. Commun. 14, 4014 (2023).

22. Khalilgharibi, N. et al. Stress relaxation in epithelial monolayers is controlled by the actomyosin cortex. Nat. Phys. 15, 839–847 (2019).

23. Duque, J. et al. Rupture strength of living cell monolayers. Nat. Mater. 23, 1563– 1574 (2024).

24. Salbreux, G., Charras, G. & Paluch, E. Actin cortex mechanics and cellular morphogenesis. Trends Cell Biol. 22, 536–545 (2012).

25. Saha, A. et al. Determining Physical Properties of the Cell Cortex. Biophys. J. 110, 1421–1429 (2016).

26. Ouzeri, A. et al. Theory of multiscale epithelial mechanics under stretch: from active gels to vertex models. bioRxiv. 2025.03.23.644792 Preprint at 10.1101/2025.03.23.644792 (2025).

27. Fischer-Friedrich, E. et al. Rheology of the Active Cell Cortex in Mitosis. Biophys. J. 111, 589–600 (2016).

28. Khoromskaia, D. & Salbreux, G. Active morphogenesis of patterned epithelial shells. eLife 12, e75878 (2023).

29. Drozdowski, O. M. & Schwarz, U. S. Morphological instability at topological defects in a three-dimensional vertex model for spherical epithelia. Phys. Rev. Res. 6, L022045 (2024).

30. Andrenšek, U., Ziherl, P. & Krajnc, M. Wrinkling Instability in Unsupported Epithelial Sheets. Phys. Rev. Lett. 130, 198401 (2023).

31. Bal, P. K., Ouzeri, A. & Arroyo, M. Continuum theory for the mechanics of curved epithelial shells by coarse-graining an ensemble of active gel cellular surfaces. bioRxiv. 2025.02.27.640501 Preprint at 10.1101/2025.02.27.640501 (2025).

32. Bico, J., Reyssat, É. & Roman, B. Elastocapillarity: When Surface Tension Deforms Elastic Solids. Annu. Rev. Fluid Mech. 50, 629–659 (2018).

33. Smeets, B., Cuvelier, M., Pešek, J. & Ramon, H. The Effect of Cortical Elasticity and Active Tension on Cell Adhesion Mechanics. Biophys. J. 116, 930–937 (2019).

34. Pogorelov, A. V. Bendings of Surfaces and Stability of Shells. (American Mathematical Soc., 1988).

35. Knoche, S. & Kierfeld, J. The secondary buckling transition: Wrinkling of buckled spherical shells. Eur. Phys. J. E 37, 62 (2014).

36. Tobasco, I. et al. Exact solutions for the wrinkle patterns of confined elastic shells. Nat. Phys. 18, 1099–1104 (2022).

37. Aharoni, H. et al. The smectic order of wrinkles. Nat. Commun. 8, 15809 (2017).

38. Biot, M. A. Folding instability of a layered viscoelastic medium under compression. Proc. R. Soc. Lond. Ser. Math. Phys. Sci. 242, 444–454 (1957).

39. Leocmach, M., Nespoulous, M., Manneville, S. & Gibaud, T. Hierarchical wrinkling in a confined permeable biogel. Sci. Adv. 1, e1500608 (2015).

40. Chatterjee, S. et al. Wrinkling and folding of thin films by viscous stress. Soft Matter 11, 1814–1827 (2015).

41. Guan, X., Nguyen, N., Cerda, E., Pocivavsek, L. & Velankar, S. S. Ridge localization driven by wrinkle packets. Soft Matter 19, 9206–9214 (2023).

42. Huang, R. & Im, S. H. Dynamics of wrinkle growth and coarsening in stressed thin films. Phys. Rev. E 74, 026214 (2006).

43. Zavodnik, J., Košmrlj, A. & Brojan, M. Rate-dependent evolution of wrinkling films due to growth on semi-infinite planar viscoelastic substrates. J. Mech. Phys. Solids 173, 105219 (2023).

44. De Simone, A., Spahr, A., Busso, C. & Gönczy, P. Uncovering the balance of forces driving microtubule aster migration in C. elegans zygotes. Nat. Commun. 9, 938 (2018).

45. Bowden, N., Brittain, S., Evans, A. G., Hutchinson, J. W. & Whitesides, G. M. Spontaneous formation of ordered structures in thin films of metals supported on an elastomeric polymer. Nature 393, 146–149 (1998).

46. Arroyo, M. & DeSimone, A. Relaxation dynamics of fluid membranes. Phys. Rev. E 79, 031915 (2009).

47. Tsuboi, A., Fujimoto, K. & Kondo, T. Spatiotemporal remodeling of extracellular matrix orients epithelial sheet folding. Sci. Adv. 9, eadh2154 (2023).

48. Tozluoǧlu, M., et al. Planar Differential Growth Rates Initiate Precise Fold Positions in Complex Epithelia. Dev. Cell 51, 299–312.e4 (2019).

49. Wyatt, T. P. J. et al. Actomyosin controls planarity and folding of epithelia in response to compression. Nat. Mater. 19, 109–117 (2020).

50. Trushko, A. et al. Buckling of an Epithelium Growing under Spherical Confinement. Dev. Cell 54, 655–668.e6 (2020).

51. Chan, C. J. et al. Hydraulic control of mammalian embryo size and cell fate. Nature 571, 112–116 (2019).

52. Prakash, V. N., Bull, M. S. & Prakash, M. Motility-induced fracture reveals a ductile-to-brittle crossover in a simple animal’s epithelia. Nat. Phys. 17, 504–511 (2021).

53. Proag, A., Monier, B. & Suzanne, M. Physical and functional cell-matrix uncoupling in a developing tissue under tension. Development 146, dev172577 (2019).

54. Sundaram, S. et al. Sacrificial capillary pumps to engineer multiscalar biological forms. Nature 636, 361–367 (2024).

