## Supplementary Information for "Multiscale wrinkling dynamics in epithelial shells"

This file includes:

- Extended figures 1-9
- Supplementary Information
    - Supplementary Movie Captions
    - Supplementary Table 1
    - Supplementary notes 1-5
    - Supplementary Refereces

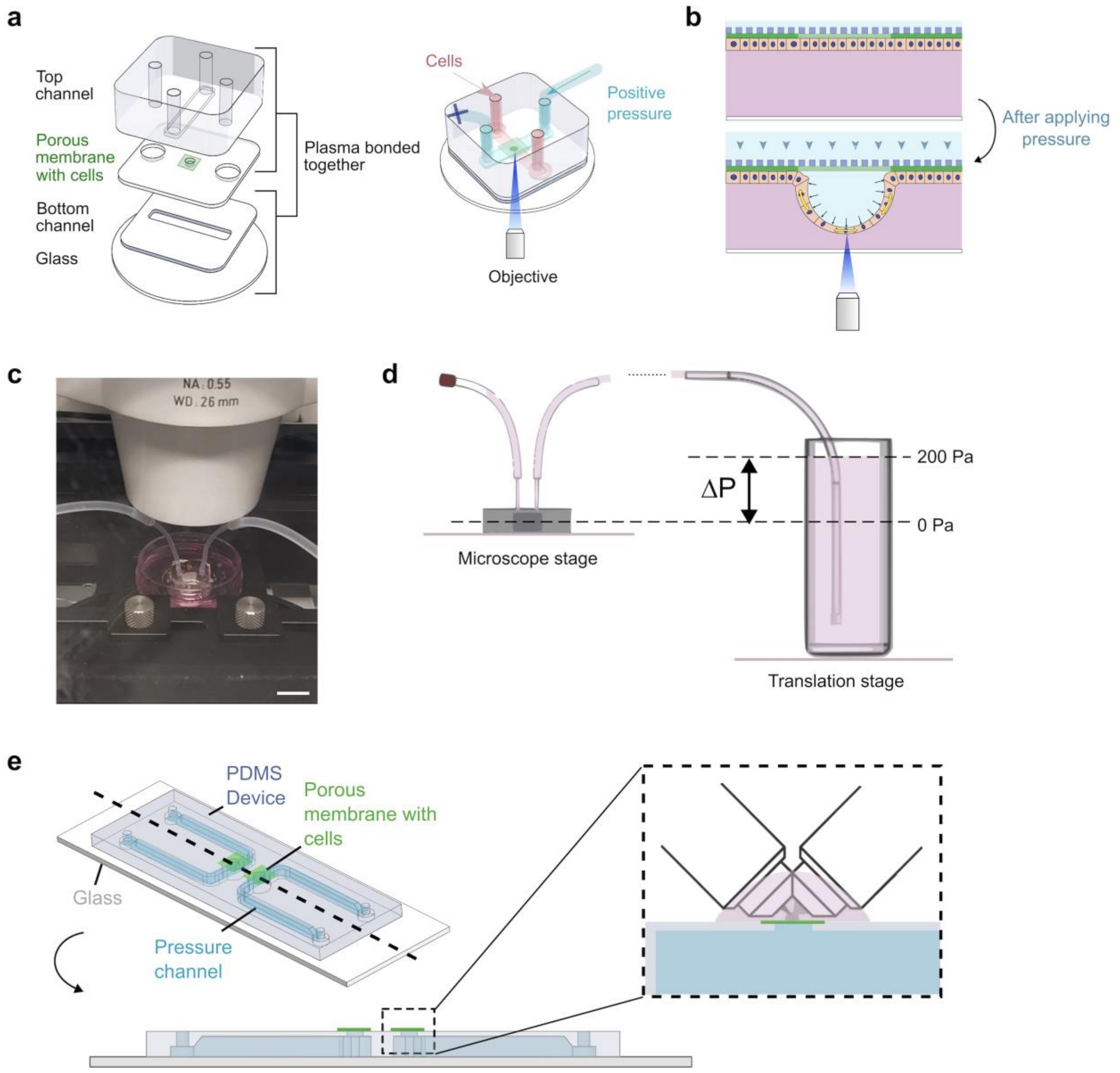


**Extended Figure 1: Microfluidic devices. a,** Schematic of four layers bonded with ozone plasma cleaning to construct MOLI. Right: Device utilized for inverted microscopy, featuring inlet and outlet channels. Only the pressure channel (blue) is connected to the tubing, with one end connected to the reservoir and the other sealed. The cell channel (red) is used for cell seeding. **b,** In this setup, the cell channel serves as the bottom channel, bringing stretched cells closer to the objective. **c**, Image of the microfluidic device on the stage of the inverted confocal microscope. Scale bar, 10 mm. **d**, The device is mounted on a microscope stage and connected to a reservoir containing medium, which is further attached to a translation stage. By adjusting the position of reservoir, hydrostatic pressure can be applied and measured. **e**, Illustration of the inflation device compatible with light sheet microscopy. The device consists of a single-piece PDMS block with two channels designed to accommodate two inflation systems on a single glass slide. The dashed line represents the cross-section of the device, and the inset indicates its use with two 40x W 0.8 NA NIR water immersion objectives at a 45-degree angle.

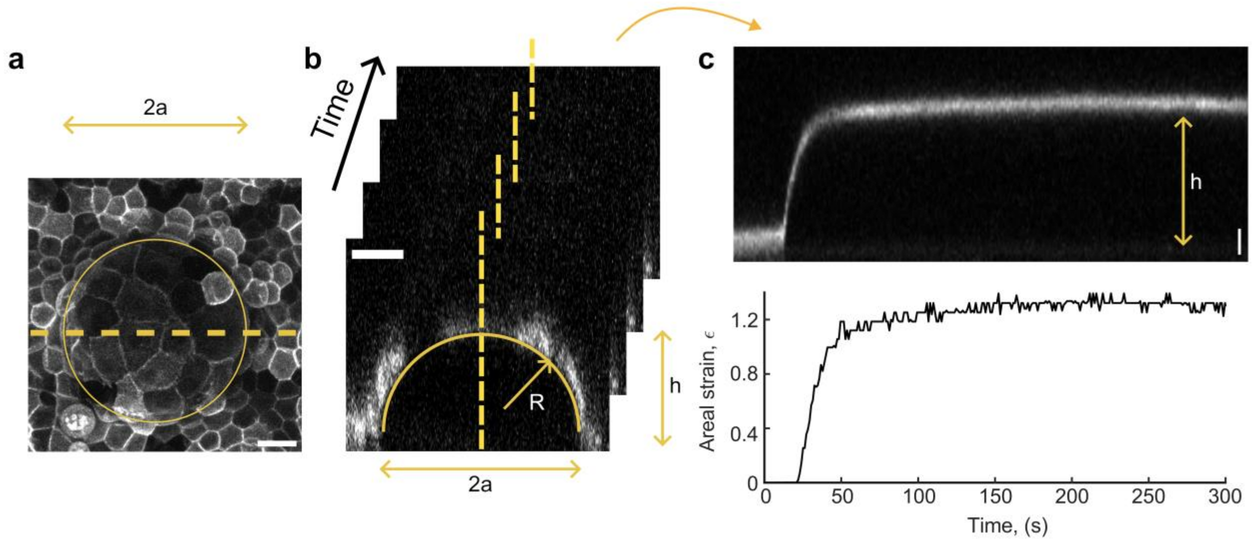


**Extended Figure 2: Quantification of shell dynamics**. **a**, Maximum intensity projection of an epithelial shell, with MDCK cells expressing CIBN-GFP-CAAX. A yellow circle marks the shell's footprint, and a dashed yellow line intersects the center of this footprint. **b**, A single XZ plane, passing through the diameter of the shell footprint, is imaged over time, enabling the measurement of the radius of curvature $R$, height $h$, and base radius $a$. **c**, A kymograph, generated from the timelapse, illustrates changes in height over time. This data can be utilized to compute areal strain $\epsilon$ over time. Scale bars, 20 μm.

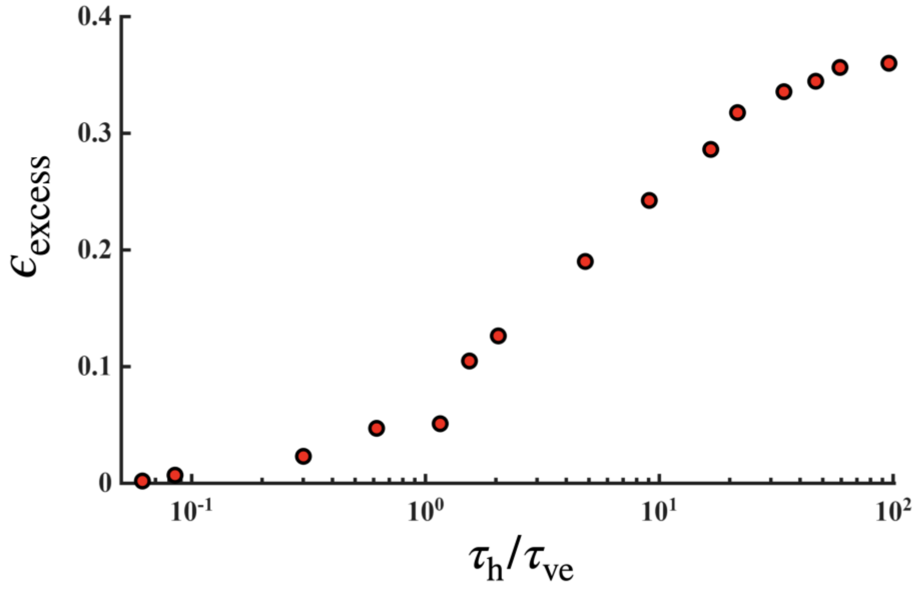


**Extended Figure 3: Excess area after full deflation as a function of the hold time according to continuum simulations**. The dimensionless excess area parameter, ϵ_excess, is defined in the main text, and the hold time is normalized by the viscoelastic relaxation time. Model parameters are reported in Supplementary Table 1, and coincide with those in Fig. 1g except for the hold time.

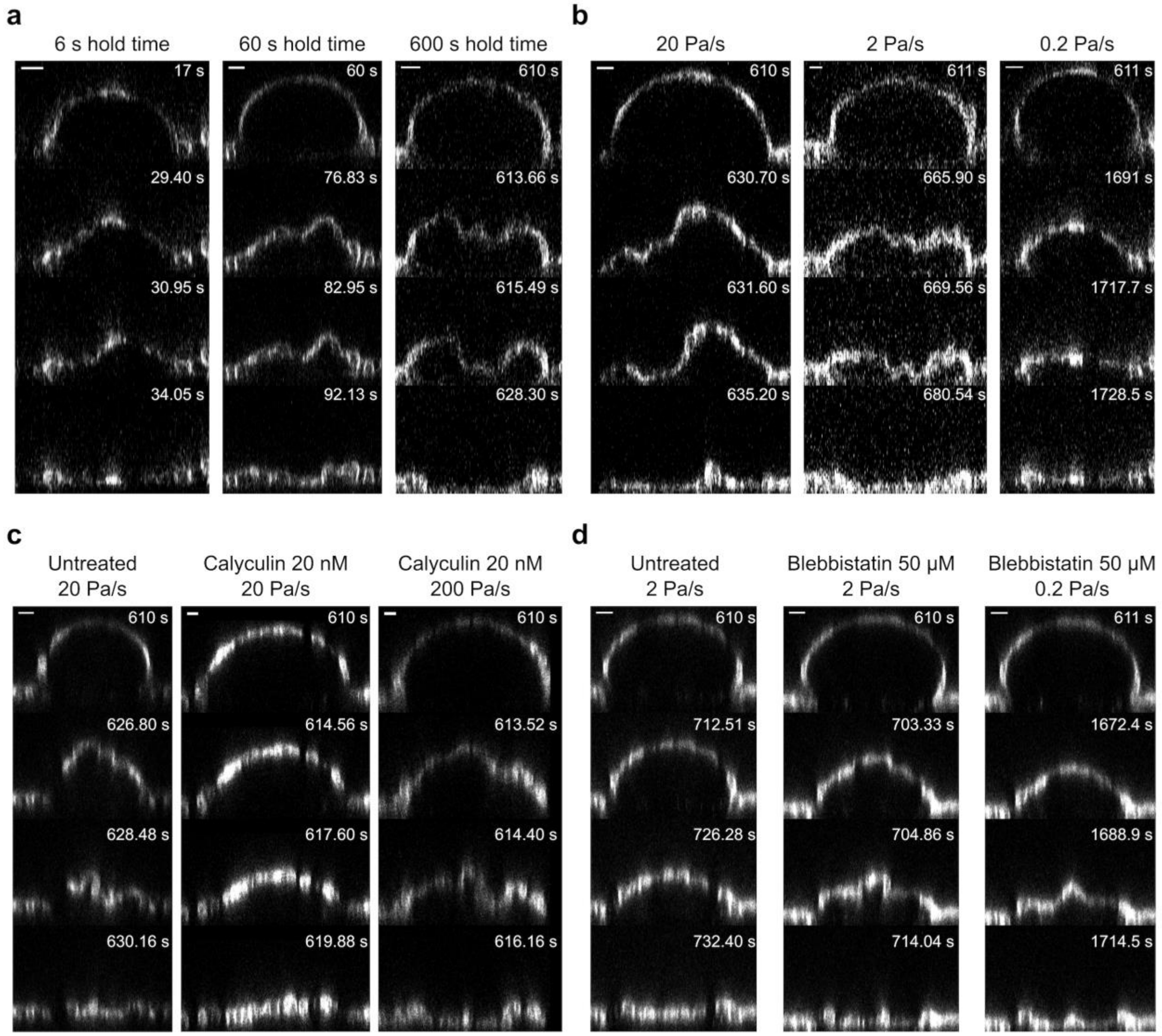


**Extended Figure 4: Representative time lapses of buckling experiments under different conditions**. **a**, Deflation of shells at a rate of 20 Pa/s, with varying hold times of 6 s, 60 s, and 600 s. **b**, Deflation of shells after a 600 s hold, with different deflation rates of 20, 2, and 0.2 Pa/s. **c**, Shells are deflated at 20 Pa/s in three conditions: untreated (left), with Calyculin 20 nM (middle), and with Calyculin 20 nM at a higher deflation rate of 200 Pa/s (right). **d**, Shells are deflated at 2 Pa/s in three conditions: untreated (left), with Blebbistatin 50 μM (middle), and with Blebbistatin 50 μM at a lower deflation rate of 2 Pa/s (right). (Scale bars, 20 μm).

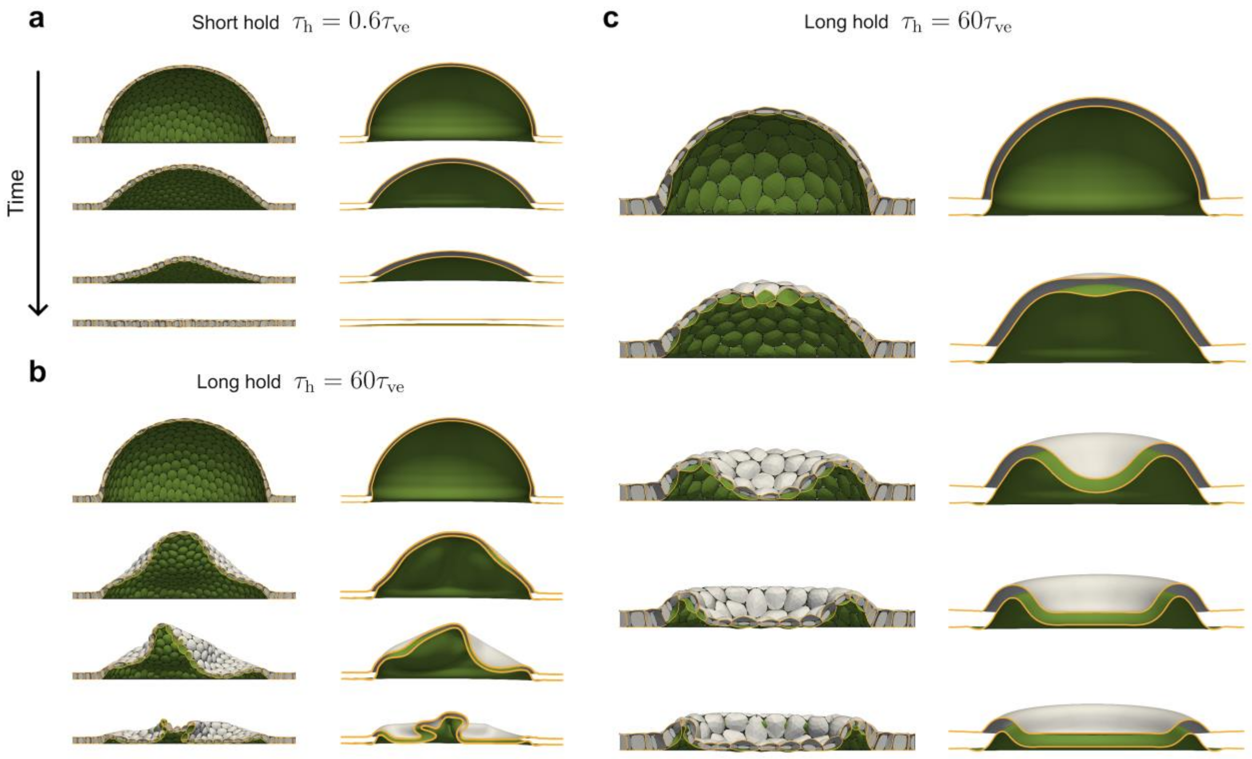


**Extended Figure 5: Comparison of the active gel vertex model and the continuum model.** Selected snapshots of rapid deflation simulations after different hold times. Panels **a** and **b** correspond to the simulations in Fig. 1g and Movie 1, whereas panel **c** corresponds to Fig. 2a. The full list of model parameters is reported in Supplementary Table 1.

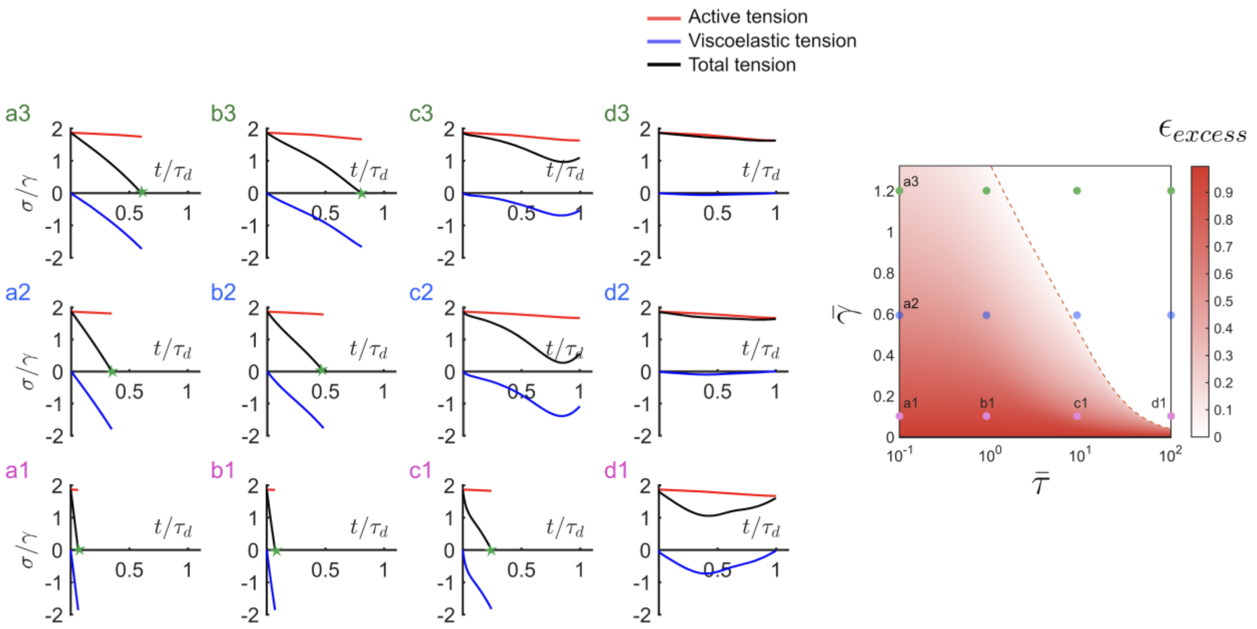


**Extended Figure 6: Time evolution of tissue tensions in different locations of the parameter space during deflation following a long hold time.** Evolution of the active, viscoelastic, and total tissue tensions $\sigma$ normalized by the cortical active tension $\gamma$ as a function of time normalized by the viscoelastic timescale. Each plot corresponds to a combination of parameters shown on the phase diagram introduced in Fig. 2c and further described in Supplementary Note 2. The green stars mark points where total tension becomes negative, leading to buckling of the epithelial shell.

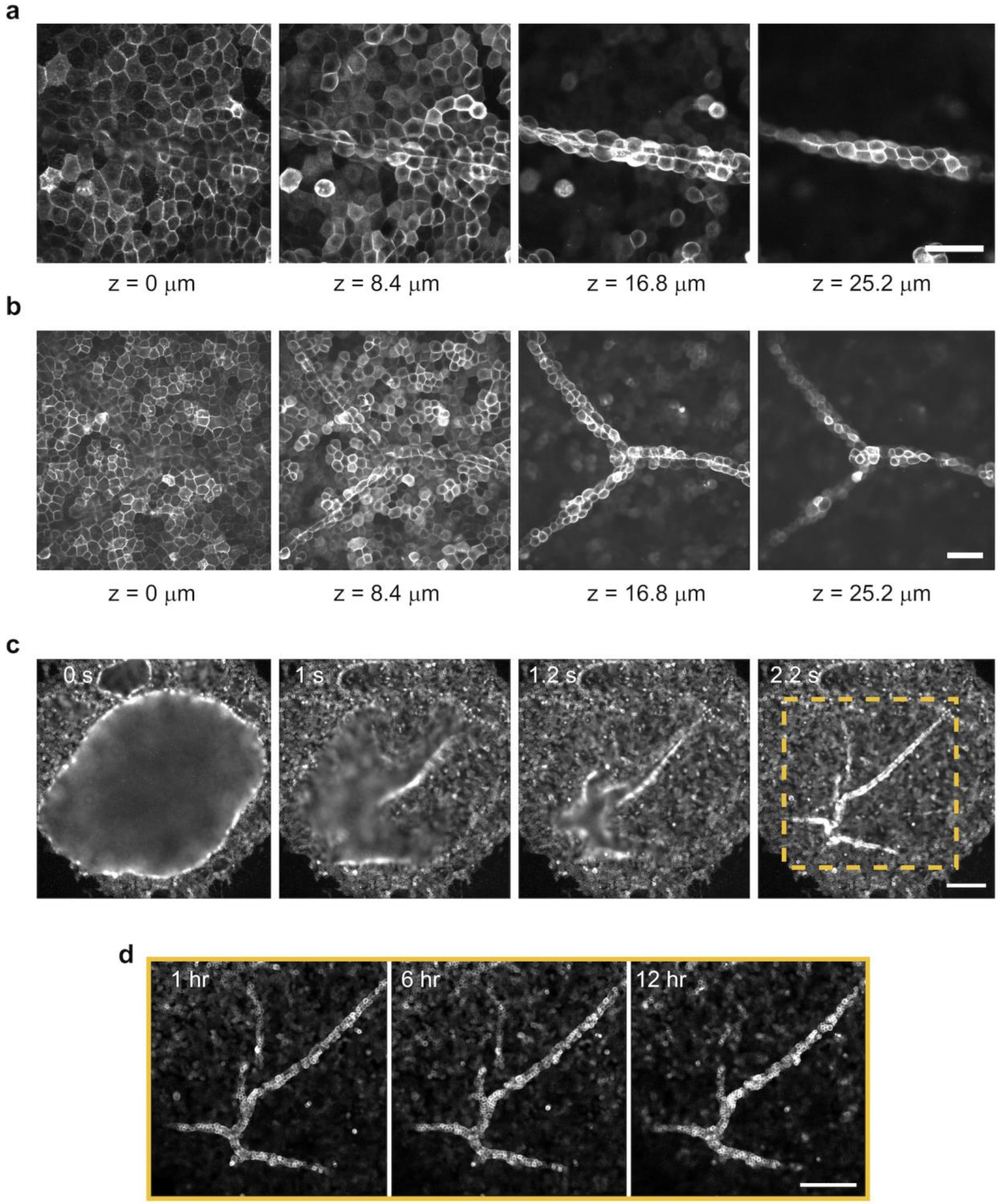


**Extended Figure 7: Post buckling patterns**. **a-b**, Confocal z-stack images reveal the wrinkles in the epithelial shells, in the case of a line (**a**) and triple junction (**b**). (Scale bars 50 µm). **c**, Epifluorescence microscopy images display the shells undergoing buckling, resulting in the formation of internal wrinkles. Scale bar, 100 µm. **d**, Timelapse of the wrinkles remaining imprinted for more than 12 hours. Scale bar, 100 µm.

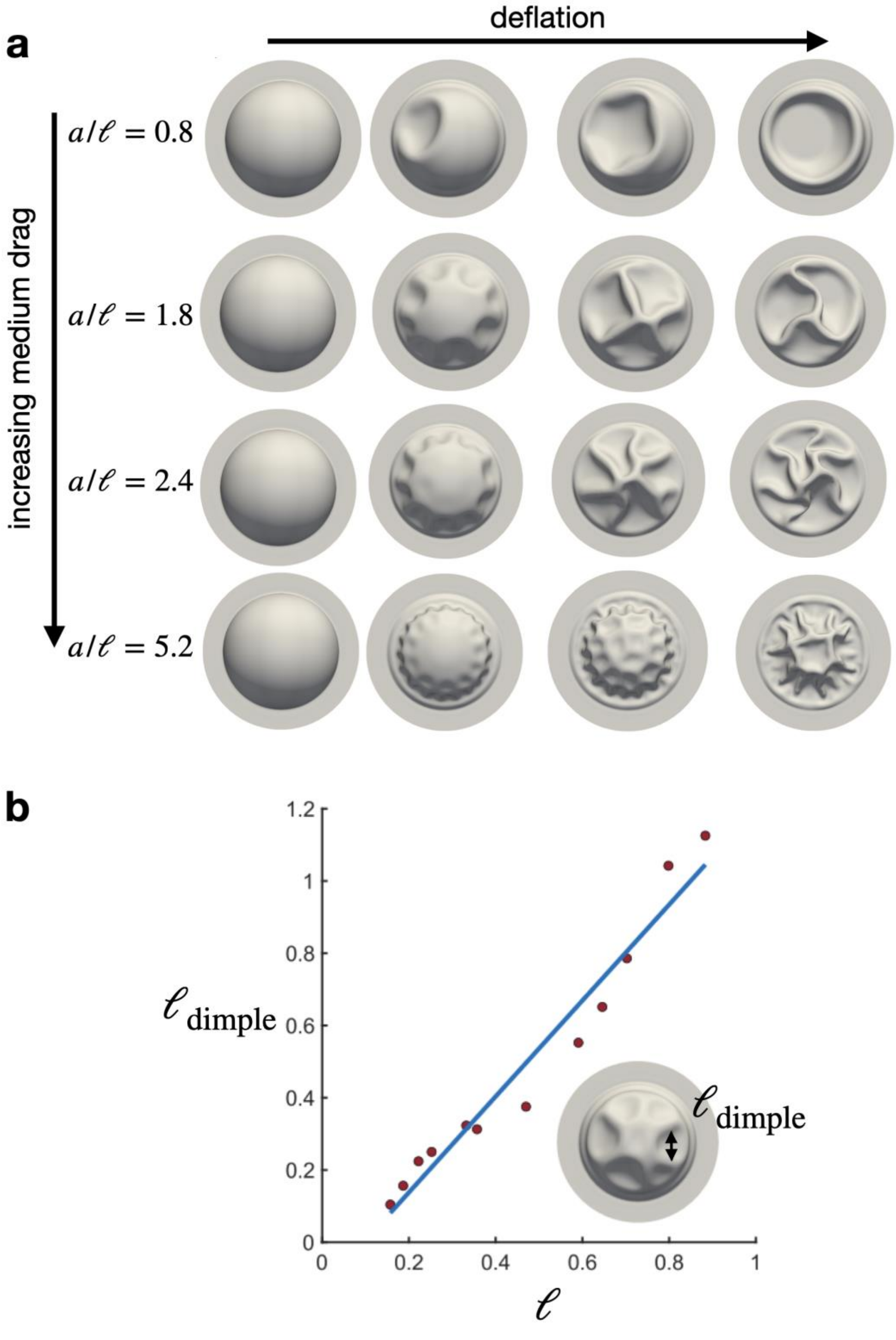


**Extended Figure 8: Continuum simulations of deflation of shells with increasing medium drag. a**, An epithelial shell of fixed slenderness $\bar{s} = 60$ is rapidly deflated after a long hold time with different magnitudes of medium drag, leading to a different ratio between the footprint radius and the dominant lengthscale of the pattern, $a/\ell$. **b**, Actual size of the dimples as measured from simulations at the onset of buckling (see inset) as a function of the predicted dominant lengthscale of the pattern $\ell$. The full list of model parameters is reported in Supplementary Table 1.

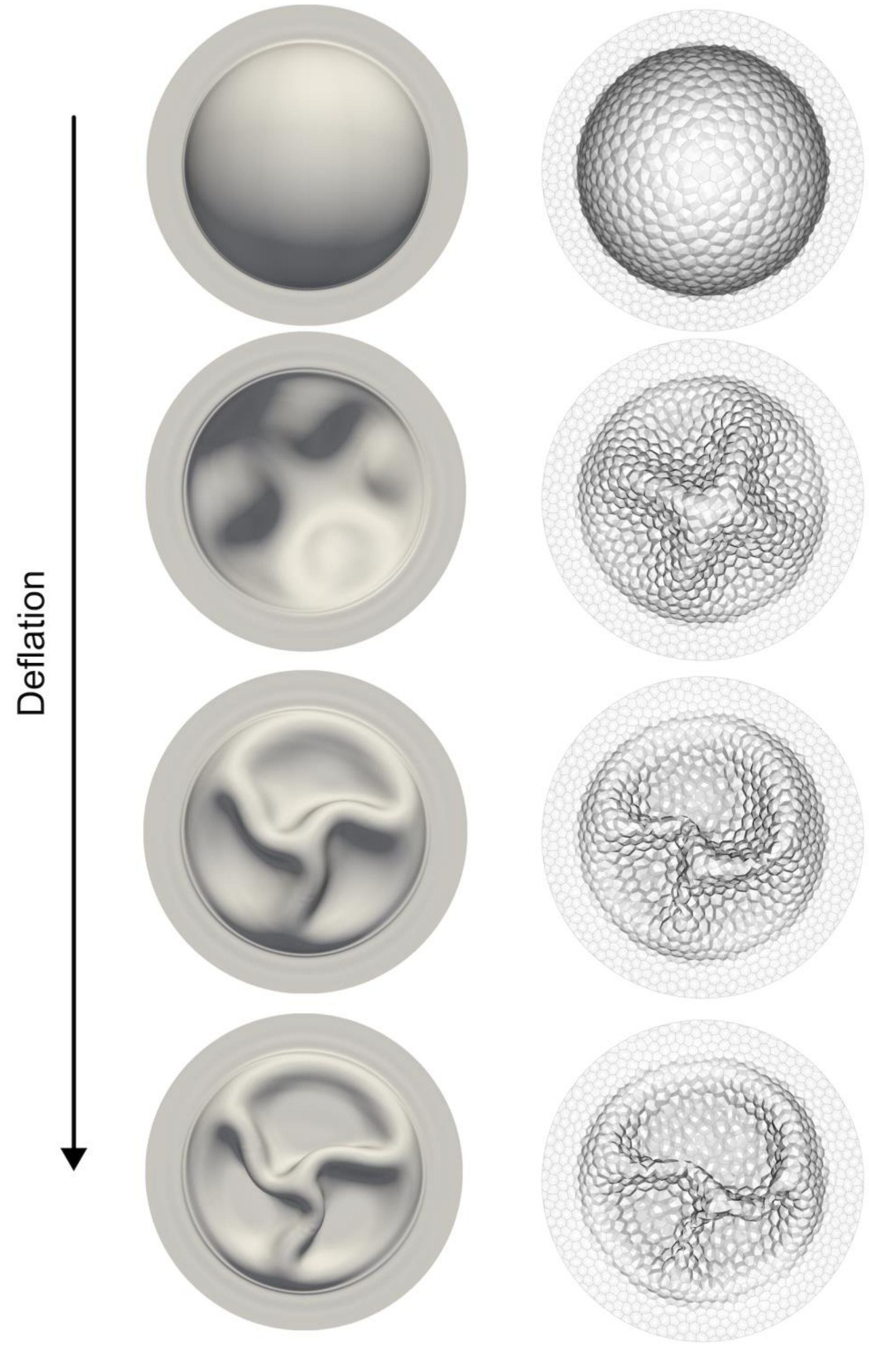


**Extended Figure 9: Comparison between the continuum simulations and those based on the active gel vertex model for a shell with internal wrinkles.** These snapshots correspond to the intermediate slenderness $\bar{s} \approx 26$ in Fig. 3c,d.

### Supplementary Information:
Multiscale wrinkling dynamics in epithelial shells

Nimesh Chahare[1,2], Adam Ouzeri[2], Thomas Wilson[1], Pradeep K. Bal[2], Tom Golde[1], Guillermo Vilanova[2], Pau Pujol-Vives[1], Pere Roca-Cusachs[1,3], Xavier Trepat[1,4,5,*], and Marino Arroyo[2,1,6,*]

[1]*Institute for Bioengineering of Catalonia (IBEC), The Barcelona Institute of Science and Technology, Barcelona, Spain*
[2]*Universitat Politècnica de Catalunya-BarcelonaTech, Barcelona, Spain*
[3]*Facultat de Medicina, Universitat de Barcelona, Barcelona, Spain*
[4]*Institució Catalana de Recerca i Estudis Avançats (ICREA), Barcelona, Spain*
[5]*Centro de Investigación Biomédica en Red en Bioingeniería, Biomateriales y Nanomedicina (CIBER-BBN), Barcelona, Spain*
[6]*Centre Internacional de Mètodes Numèrics en Enginyeria (CIMNE), Barcelona, Spain*

#### Contents

**Supplementary Movie Captions** **2**

**Supplementary Table 1: Model parameters in dimensionless form** **3**

**Supplementary Note 1: Computational modeling of epithelial shells** **4**

**Supplementary Note 2: Building the phase diagram** **6**

**Supplementary Note 3: Selection of material parameters** **7**

**Supplementary Note 4: Scaling argument for the wrinkling wavelength** **8**

**Supplementary Note 5: Model for sub-cellular wrinkling** **11**

**Supplementary References** **12**

#### Supplementary Movie Captions

**Movie 1:** Comparison between the active gel vertex model and the continuum model undergoing rapid deflation after different hold times.

**Movie 2:** Representative 3D light sheet microscopy data of a shell undergoing bucking, with two states: unbuckled (left) and buckled (right). Scale bar 20 µm.

**Movie 3:** Representative line scan microscopy videos of mid-section of shells deflating at 20 Pa/s after hold times of 6, 60, and 600 seconds, and shells deflating at rates of 0.2, 2, and 20 Pa/s with a hold time of 600 seconds. Scale bar: 20 µm.

**Movie 4:** Representative line scan microscopy video of mid-section of shells deflating with drug treatments of Calyculin A 20nM and Blebbistatin 50 µM. Scale bar 20 µm.

**Movie 5:** Representative confocal microscopy video at the base of shells of different sizes. Scale bar 100 µm.

**Movie 6:** Continuum simulation of a shell exhibiting a polygonal dimple.

**Movie 7:** Continuum simulations of deflating shells with increasing medium drag.

**Movie 8:** Continuum simulations of deflating shells with increasing size.

**Movie 9:** Simulations of equibiaxial and uniaxial compression of a tissue modeled by an active gel vertex model explicitly accounting for cytosolic viscous hydrodynamics and cortical bending rigidity.

**Movie 10:** Mean curvature of stretched and buckled cell.

**Movie 11:** Representative confocal microscopy video at the base of shells of different shapes. Scale bar 100 µm.

**Movie 12:** Continuum simulations of deflating shells of different shapes.

**Supplementary Table 1: Model parameters in dimensionless form.** Symbols are defined in the main text and in Supplementary Note 1. Parameter $\tau_{to}/\tau_{ve}$ only affects simulations based on the active gel vertex model, since we simplify turnover dynamics in the continuum theory as explained above.

| | $\bar{\gamma}$ | $\tau_{to}/\tau_{ve}$ | $\bar{f}$ | $\bar{s} = 2a/h_0$ | $\tau_h/\tau_{ve}$ | $\bar{\tau}$ | $a/\ell$ |
|---|---|---|---|---|---|---|---|
| Fig. 1g, Ext. Fig. 5a,b, Movie 1 | 0.5 | 2 | 0.19 | 26 | 0.6/60 | 0.6 | 1.5 |
| Fig. 2a, Ext. Fig. 5c | 0.5 | 2 | 0.19 | 13 | 60 | 0.6 | 0.75 |
| Ext. Fig. 3 | 0.5 | - | 0.19 | 26 | 0.01 to 100 | 0.6 | 1.5 |
| Movie 6 | 0.5 | - | 0.19 | 60 | 60 | 0.6 | 0.8 |
| Ext. Fig. 8, Movie 7 | 0.5 | - | 0.19 | 60 | 60 | 0.6 | 0.8 to 5.2 |
| Fig. 3c-e, Movie 8 | 0.5 | 2 | 0.19 | 13, 26, 70 | 60 | 0.6 | 0.75, 1.5, 4 |
| Ext. Fig. 9 | 0.5 | 2 | 0.19 | 26 | 60 | 0.6 | 1.5 |
| Fig. 5c, Movie 12 | 0.5 | - | 0.19 | ~ 70 | 60 | 0.6 | ~ 4 |

### Supplementary Note 1: Computational modeling of epithelial shells

##### 3D active gel vertex model

To capture the hierarchical mechanical organization of epithelial shells and the dominant role of the actomyosin cytoskeleton in determining the mechanical properties of cell monolayers under stretch[20;23;49], we first turned to a model explicitly describing each individual cell. In this model, the mechanical behavior of each cellular surface is described with an active gel model of the actomyosin cortex capturing its viscoelasticity, active contractility and turnover. In this model, akin to a 3D vertex model with curved cellular surfaces, each cell surface is triangulated and the active gel model is solved on it using finite elements. Cellular surfaces enclose fixed cellular volumes. We include a repulsive potential between the surfaces of different cells to preclude their interpenetration. The active gel model and this approach are described in detail elsewhere[26].

##### Continuum theory for epithelial shells

We complemented this 3D active gel vertex model with a fully nonlinear continuum theory for epithelial shells. This theory coarse-grains the 3D cellular model by relating the deformation of apical, basal and lateral surfaces of cells to the continuum deformation of the tissue mid-surface and a thickness director field. In this model, the ensemble of apical and basal surfaces are modeled as offset surfaces to the shell mid-surface. The mechanical effect of lateral surfaces is also accounted for. It retains the active gel description of cellular surfaces. It is described in detail elsewhere[31]. The continuum model affords a significant reduction of the computational complexity, enabling much faster simulations. Simulations presented in the present work (Figs. 2a, 3c-d, Extended Figs. 5 and 9 and Movie 1) and in this reference demonstrate that this continuum shell theory mimics very accurately the 3D active gel vertex model. The continuum theory is not applicable in its present form when the cell monolayer exhibits significant rearrangements of the junctional network, or when cellular deformations are not affine, two situations not observed in our experiments.

##### Model parameters

We first describe the material parameters of the active gel. This model tracks the dynamics of the cortical density or thickness, that evolves due to deformation and turnover, and the dynamics of material metric tensors, which model the evolution of the reference configuration of the material as it relaxes viscoelastically due to network remodeling. This viscoelastic relaxation tends to lower the stored elastic energy and is driven by elastic stress in the network. The physical properties of the cortex (elasticity, viscosity, active tension) are assumed to be proportional to the cortex density. The short-term elasticity of the cell cortex is characterized by the Lamé elastic parameters $\lambda$ (dilatational) and $\mu$ (shear). We assume that $\lambda = \mu$[55]. These parameters describe reference values at the steady-state density, and represent integrated elastic moduli through the thickness, and therefore have units of surface tension. The active contractile tension for the reference density is $\gamma_a$. The surface viscosity parameter for the reference density, $\eta$, controls the viscoelastic relaxation. We can thus define the viscoelastic time-scale as $\tau_{ve} = \eta/\mu$. The turnover time-scale controlling density dynamics is denoted as $\tau_{to}$. These parameters allow us to define several dimensionless quantities, notably the ratio of turnover to viscoelastic times $\tau_{to}/\tau_{ve}$, and the elastocapillary parameter $\bar{\gamma} = \gamma_a/\mu$.

Our models view tissues as collections of active gel surfaces enclosing constant volumes. The interplay between apical, basal and lateral properties of these active gels controls the mechanical properties of the tissue. Here, we assume that apical and basal surfaces have the same mechanical properties, since we have shown earlier that apicobasal asymmetries have little impact on the mechanics of inflated epithelial monolayers[21]. However, we allow lateral surfaces to have different properties, and assume that all mechanical properties of lateral surfaces ($\mu$, $\eta$ and $\gamma$) are scaled by the same factor with respect to apicobasal properties. We denote by $\gamma_a^{ab}$ and $\gamma_a^{lat}$ the apicobasal and lateral active tensions. As identified in[26;31], the cell aspect ratio and the ratio of lateral to apicobasal active tensions combine into a dimensionless prestress parameter defined as

$$\bar{f} = f_0 \frac{\gamma_{\mathrm{a}}^{\mathrm{lat}}}{\gamma_{\mathrm{a}}^{\mathrm{ab}}}, \tag{1}$$

where $f_0$ is the specific lateral area in a given state of the tissue, that is the area of lateral surfaces per unit tissue area. For an idealized crystalline tissue with hexagonal cells of apical/basal area $S_0$ and height $h_0$, we can explicitly evaluate $f_0$ leading to

$$\bar{f} = \sqrt{\frac{\sqrt{3}}{8}} \frac{h_0}{\sqrt{S_0}} \frac{\gamma_{\mathrm{a}}^{\mathrm{lat}}}{\gamma_{\mathrm{a}}^{\mathrm{ab}}} \approx 0.46 \frac{h_0}{\sqrt{S_0}} \frac{\gamma_{\mathrm{a}}^{\mathrm{lat}}}{\gamma_{\mathrm{a}}^{\mathrm{ab}}}. \tag{2}$$

When $\bar{f} > 1$, then the tissue is under compression in its steady state (when all cortical elastic stresses have dissipated). Conversely, when $\bar{f} < 1$, then the tissue is in tension in its steady state. This parameter needs to be defined at a specific configuration. Here, we define it in the planar state of the tissue prior to inflation.

Our simulations with the 3D active gel vertex model show that the main tension dynamics in the tissue are the result of viscoelastic relaxation, with a relatively small effect of density variations due to stretch and turnover. Accordingly, we simplify our simulations based on the continuum theory of epithelial shells by assuming that the cortex is always at its steady-state density, which reduces the number of equations to be solved and speeds up simulations[31].

As discussed in the main text, we further define dimensionless parameters associated with the slenderness of the epithelial shells, $\bar{s} = 2a/h_0$ (where $a$ is the radius of the footprint), and with the hold and deflation times relative to the viscoelastic time, $\tau_{\mathrm{h}}/\tau_{\mathrm{ve}}$ and $\bar{\tau} = \tau_{\mathrm{d}}/\tau_{\mathrm{ve}}$.

Finally, we incorporate the effect of the viscous drag by the fluid medium of viscosity $\eta_{\mathrm{m}}$ embedding the tissue, which introduces the dimensionless number $\eta_{\mathrm{m}} h_0/\eta$. An accurate description of this effect involves solving the 3D Stokes flow around the deforming epithelial shell. To reduce computational cost, here we simplified this effect as a local viscous drag, such that each point in the epithelial shell experiences a drag force density per unit area of the form

$$\boldsymbol{f}_{\mathrm{drag}} = -\tilde{\eta}_{\mathrm{m}} \boldsymbol{v}, \tag{3}$$

where $\tilde{\eta}_{\mathrm{m}}$ is a viscous drag coefficient and $\boldsymbol{v}$ is the local velocity of the shell at that point. As discussed in the main text, the main role of the viscous drag is to control the wrinkling wavelength of the tissue $\ell$, given by Eq. (7) or Eq. (18) depending on the model for viscous drag. Under the local viscous drag approximation of our simulations, $\ell$ is given by Eq. (18). We report the viscous drag coefficient $\tilde{\eta}_{\mathrm{m}}$ in dimensionless form using the parameter $a/\ell$.

Supplementary Table 1 reports the model parameters in dimensionless form for each simulation in the paper, except for those in Fig. 4a-c and Movie 9, which are reported in Supplementary Note 5. Our choices are justified in Supplementary Note 3.

#### Supplementary Note 2: Building the phase diagram

To build the phase diagram in Fig. 2c, we analyze a tissue undergoing deflation and check whether the condition $\sigma_t < 0$ is met prior to full deflation, where $\sigma_t = \sigma_a + \sigma_{ve}$ is the total tissue tension, composed of an active and a viscoelastic component. Epithelial shells being quite slender, this condition should approximate well the buckling condition (see also the discussion following Eq. (10)). We make the approximation that strain in a deflating epithelial shell is uniform and isotropic, which is quite accurate except close to the boundaries. Further neglecting the effect of curvature, we can analyze a planar and uniform tissue, for we have explicit expressions for the tension[26;31]. The evaluation of tension requires integrating ordinary differential equations for the material metric tensors capturing viscoelastic relaxation, which is computationally straightforward. Fixing the parameters $\bar{f} = 0.19$ and $\tau_h/\tau_{ve} = 60$, we sweep $\bar{\gamma}$ and $\bar{\tau}$. For each parameter combination, starting from an areal strain of $\epsilon = 1$ (corresponding to a hemispherical shell) and after a long hold time, we compress the tissue and record whether the condition $\sigma_t < 0$ is met or not before $\epsilon = 0$ (the planar state). When buckling takes place, we estimate the normalized excess area $\epsilon_{excess} = A/(\pi a^2) - 1$ as the areal strain when the condition $\sigma_t = 0$ is met. This approximation is justified by our 3D nonlinear simulations, which show that the tissue area changes very little after buckling. To test the accuracy of this approach to build the buckling phase diagram, we performed 20 nonlinear epithelial shell simulations across the parameter regime, finding excellent agreement.

#### Supplementary Note 3: Selection of material parameters

To estimate the prestress parameter $\bar{f}$, other reports[20;21] show that our tissues are in state of positive tension prior to inflation, with a relatively flat shape of the relation between tissue tension and areal strain. This indicates that $\bar{f} < 1$ and that this parameter is significantly smaller than 1, Supplementary Fig. 1 and references[26;31]. Accordingly, we choose in all of our calculations $\bar{f} \approx 0.2$.

Our experiments show that the critical hold time for buckling is of about 60 s, Fig. 1i. Based on simulations, Extended Fig. 3 shows that the critical hold time is between 1 and 10 times $\tau_{ve}$, and hence we can estimate $\tau_{ve} \approx 10$ s, which agrees with previous measurements in tissues and single cells[22;25;27]. To estimate $\bar{\gamma}$, we estimated the threshold deflation time for buckling in our experiments as 100 s, Fig. 2i, i.e. $\bar{\tau}^* \approx 10$. Being a threshold value, the pair $(\bar{\tau}^*, \bar{\gamma})$, should lie on the dotted line in the phase diagram in Fig. 2c. We can thus estimate $\bar{\gamma} \approx 0.5$. This value is comparable to the low-frequency value from single-cell rheology[27].

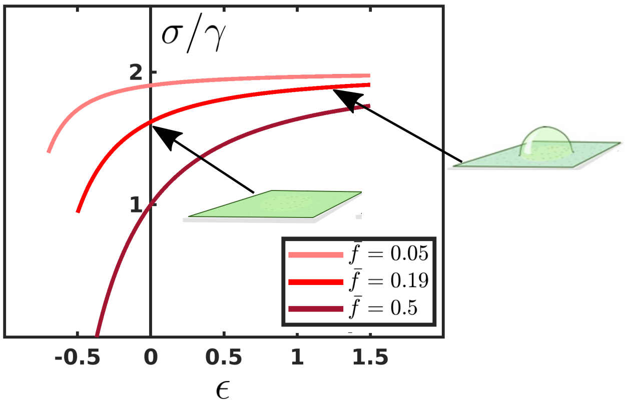


**Supplementary Figure 1:** Tissue tension normalized by $\gamma_a^{ab}$ at steady-state as a function of the areal strain for different values of the prestress parameter $\bar{f}$.

### Supplementary Note 4: Scaling argument for the wrinkling wavelength

Compressed thin films mechanically confined by a surrounding medium develop wrinkling patterns, whose wavelength depends on the properties of the thin film and of the surrounding medium. In a pioneering work in this field, Biot studied the problem in a rather general setting[38], predicting the dominant wrinkling wavelength and critical load considering a viscoelastic layer surrounded by a viscoelastic medium, with purely elastic or viscous materials as limiting cases. For an elastic thin film on a surrounding elastic medium, the dominant wavelength is set by a competition between elastic energy in the thin film, favoring long wavelengths, and elastic energy in the surrounding medium, favoring short wavelengths. If the elastic thin film is surrounded by a viscous fluid or a viscoelastic medium, then the energy dissipation in the surrounding medium as the wrinkling instability dynamically develops or as the film is dynamically loaded also affects the wavelength selection. In this more complex situation, more recent work in viscous fluids[39;40;41 56;57] or viscoelastic media[58 42;43] shows that the dominant wavelength depends on the loading protocol (stress or strain controlled, step or ramp application) and time, as the wrinkle pattern often coarsens into folds.

In our system, epithelial shells are rapidly deflated, which results in compression of the cell monolayer and the cortical surfaces within. Because epithelial shells are surrounded by a viscous fluid and the cortical surfaces are surrounded by the more viscous cytosol, the wrinkling pattern at the onset of buckling of these structures could result from the interplay between thin film mechanics and viscous stresses in the surrounding medium. A detailed analysis of our system is difficult given the geometry, the active viscoelastic behavior of the cell cortex and the tissue, and the loading protocol. Instead, we provide a rough estimate of the dominant wavelength using dimensional analysis and several simplifying assumptions. We first assume that the deflation time is short compared to the viscoelastic relaxation time ($\bar{\tau}$ small), so that $\tau_d$ is the only important time-scale in the system. We thus ignore for this analysis the viscoelastic relaxation taking place while the instability develops. Because wavelength selection is the result of a competition between bending and viscous dissipation, controlled by the bending rigidity $D$, the viscosity of the surrounding medium $\eta_m$ and the strain rate, we can only form one length-scale

$$\ell = \sqrt[3]{\frac{D}{\eta_m \dot{\epsilon}}}, \tag{4}$$

where $\eta_m$ is the viscosity of the surrounding medium and $\dot{\epsilon}$ is the strain rate. This expression agrees up to prefactors with that by Biot[38]. The dominant wavelength at the onset of buckling, before pattern coarsening takes place in the fully nonlinear post-buckling regime, should be commensurate to this lengthscale. This estimation may have several limitations, such as the fact that other dimensionless parameters in the model such as $\bar{\gamma}$ and $\bar{\tau}$ could play a role, the effect of spherical geometry, which introduces an additional lengthscale, or the underestimation of the drag caused by the fluid in the confined lumen under the epithelial monolayer.

A similar scaling argument shows that the critical compressive tension for buckling should scale as

$$\sigma^{\text{crit}} = -D^{1/3} (\eta_m \dot{\epsilon})^{2/3} . \tag{5}$$

Since we start deflation from a strain of approximately 1 (a hemisphere), we can estimate the strain rate as $\dot{\epsilon} \approx 1/\tau_d$.

#### Cell monolayer

Since at fast loading rates the cortex behaves as an elastic material with modulus $\mu$, and since this elastic material resisting bending is essentially concentrated at the apical and basal surfaces separated by a distance $\sim h_0$, see reference[31], the bending rigidity of the cell monolayer can be estimated as

$$D \approx \frac{\mu h_0^2}{2}. \tag{6}$$

The dominant wavelength at the onset of buckling then takes the form

$$\ell_{\text{tissue}} = \sqrt[3]{\frac{\mu h_0^2 \tau_d}{2\eta_m}} = h_0 \sqrt[3]{\frac{\mu \tau_d}{2 h_0 \eta_m}} = h_0 \sqrt[3]{\frac{\bar{\tau}}{2} \frac{\eta}{h_0 \eta_m}}. \tag{7}$$

In the second expression, the inside the cubic root is a dimensionless parameter comparing the stiffness of the thin sheet to an effective stiffness of the surrounding medium. In the case of a very thin film of surrounding fluid described using lubrication theory, a different scaling is obtained, but this dimensionless number is shown to play a central role in the physics of viscous wrinkling[41]. The third expression expresses $\ell_{\text{tissue}}/h_0$ in terms of the dimensionless deflation rate and the dimensionless number comparing bulk and cortical viscosity.

We expect the transition between peripheral folds and internal folds to take place when the system size $2a \approx \ell_{\text{tissue}}$, which defines a critical slenderness for this transition given by

$$\bar{s}_{\text{crit}} \approx \sqrt[3]{\frac{\mu \tau_d}{2 h_0 \eta_m}} = \sqrt[3]{\frac{\bar{\tau}}{2} \frac{\eta}{h_0 \eta_m}}. \tag{8}$$

Tension in the tissue can be approximated as twice the tension in each cortical surface, which for a prestressed-elastic behavior takes the form $\sigma \approx 2(\gamma + \mu\epsilon)$. This leads to the following estimate for the critical strain at buckling

$$\epsilon_{\text{tissue}}^{\text{crit}} \approx -\left[\bar{\gamma} + \frac{1}{2^{4/3}} \left(\frac{h_0 \eta_m}{\mu \tau_d}\right)^{\frac{2}{3}}\right]. \tag{9}$$

Since $\bar{\gamma}$ is comparable to 1, we take an elastic modulus comparable to cortical tension, $\mu \approx 1$ mN/m. As for the strain rate, taking $\tau_{ve} \approx 10$ s and $\bar{\tau} = 0.1$, we estimate $\dot{\epsilon} \approx 1\ \text{s}^{-1}$. We approximate the viscosity of the medium as that of water, $\eta_m \approx 10^{-3}$ Pa s and the monolayer thickness as $t_0 \approx 10$ µm. We then estimate

$$\ell_{\text{tissue}} \approx 370\ \mu\text{m}, \qquad \bar{s}_{\text{crit}} \approx 37, \qquad \epsilon_{\text{tissue}}^{\text{crit}} \approx -\left[\bar{\gamma} + 2 \cdot 10^{-4}\right]. \tag{10}$$

The third estimation shows that the critical compressive elastic tension is essentially that needed to cancel the positive active tension, as assumed to generate the phase diagram in Fig. 2b.

As discussed in the main text, the key parameter that controls the number of internal dimples at buckling onset and the final number of wrinkles is $a/\ell_{\text{tissue}}$. Experimentally, we change this parameter by a factor of 5.3 by changing $a$ by the same factor. If we wanted to achieve a similar factor by changing viscosity, Eq. (7) shows that we would need to increase $\eta_m$ by a factor of 150.

#### Cortical surface

An analogous argument can be applied to a cortical surface of thickness $h_{\text{cortex}}$, whose bending rigidity can be estimated as

$$D \approx \frac{\mu h_{\text{cortex}}^2}{12}, \tag{11}$$

leading to

$$\ell_{\text{cortex}} = h_{\text{cortex}} \sqrt[3]{\frac{\mu \tau_d}{12 h_{\text{cortex}} \eta_c}}, \tag{12}$$

with $\eta_c$ the viscosity of the cytosol, which is significantly larger than that of the surrounding medium. The critical slenderness now takes the form

$$\bar{s}_{\text{crit}} \approx \sqrt[3]{\frac{\mu \tau_d}{12 h_{\text{cortex}} \eta_c}}. \tag{13}$$

We can also estimate

$$\epsilon_{\mathrm{cortex}}^{\mathrm{crit}} \approx -\left[\bar{\gamma} + \frac{1}{12^{1/3}}\left(\frac{h_{\mathrm{cortex}}\eta_{\mathrm{c}}}{\mu\tau_{\mathrm{d}}}\right)^{\frac{2}{3}}\right]. \tag{14}$$

Using the estimation $\eta_{\mathrm{c}} \approx 6 \cdot 10^{-1}$ Pa s for the cytosolic viscosity[44] and $h_{\mathrm{cortex}} \approx 100$ nm for the cortical thickness, we obtain

$$\ell_{\mathrm{cortex}} \approx 1\ \mu\mathrm{m}, \qquad \bar{s}_{\mathrm{crit}} \approx 11, \qquad \epsilon_{\mathrm{cortex}}^{\mathrm{crit}} \approx -\left[\bar{\gamma} + 7 \cdot 10^{-4}\right]. \tag{15}$$

More importantly, just requiring an estimation of the ratios of cortical to tissue thickness and of cytosol to medium viscosity, we have

$$\frac{\ell_{\mathrm{cortex}}}{\ell_{\mathrm{tissue}}} \approx \frac{1}{\sqrt[3]{6}}\left(\frac{h_{\mathrm{cortex}}}{h_0}\right)^{\frac{2}{3}}\left(\frac{\eta_{\mathrm{m}}}{\eta_{\mathrm{c}}}\right)^{\frac{1}{3}} \approx 3 \cdot 10^{-3}. \tag{16}$$

#### Simulations with approximate model for viscous drag

In our simulations, we do not resolve the viscous hydrodynamics of the embedding medium. Instead, we approximate this effect using a local viscous drag according Eq. (3) with drag coefficient $\tilde{\eta}_{\mathrm{m}}$. With this model, the scaling of the dominant wavelength becomes

$$\ell_{\mathrm{tissue}} = \sqrt[4]{\frac{D}{\tilde{\eta}_{\mathrm{m}}\dot{\epsilon}}}, \tag{17}$$

which for a cell monolayer deflating at strain rate $\dot{\epsilon} = 1/\tau_{\mathrm{d}}$ can be approximated as

$$\ell_{\mathrm{tissue}} \approx h_0 \sqrt[4]{\frac{\mu\tau_{\mathrm{d}}}{2h_0^2\tilde{\eta}_{\mathrm{m}}}} = h_0 \sqrt[4]{\frac{\bar{\tau}}{2}\frac{\eta}{h_0^2\tilde{\eta}_{\mathrm{m}}}}. \tag{18}$$

However, this simplification does not change the fact that drag should result in internal folds when system size becomes larger than the dominant wavelength. Despite all the approximations made to reach this expression, our numerical simulations show that the wavelength of the pattern of dimples at the onset of buckling agrees very well with this estimation, Extended Fig. 8.

### Supplementary Note 5: Computational model for sub-cellular wrinkling

To describe sub-cellular wrinkling, we had to further enrich the 3D active gel vertex model to account for the hydrodynamics of the cytosol and the bending rigidity of cortical surfaces. The local approximation in Eq. (3) cannot be used for the cortical surfaces if we want to be able to describe the coexistence of tissue and cortical buckling.

To address this, we discretized the interior of cells with a tetrahedral finite element mesh, conforming with the surface mesh of cortical surfaces. In this bulk mesh, we solved the Stokes equations for the low Reynolds number hydrodynamics of a Newtonian fluid. We used a Lagrangian description of the fluid, which given the highly constrained cytosolic flows does not lead to severe mesh distortion and simplifies the implementation. We ignored a more complex rheology of the cytosol or the presence of large organelles such as the nucleus. To account for bending rigidity, we implemented a Helfrich model and used subdivision surface finite elements, a Spline technique supported on surface triangulation that allows us to compute the curvature directly[59]. See Supplementary Figure 2 for an illustration.

Solving an entire spherical epithelial shell with this model is computationally expensive. Instead, we consider a small tissue composed of a few cells in a very stretched configuration, corresponding to a selection of cells in an inflated shell, and then rapidly compress this system. See Movie 9 for an illustration of biaxial and uniaxial compression.

In the simulations of Fig. 4a-c, we consider a high strain rate placing us in the small $\bar{\tau}$ limit, and consider $\bar{\gamma} = 0.25$. We choose $\gamma_a^{lat}/\gamma_a^{ab} = 0.4$. Since we model compression starting from a highly stretched state representative of an inflated shell, we choose the ratio between the cell thickness $h_0$ and the radius $R$ of hexagon modeling the apical surface to be small, $h_0/R = 0.2$. This leads to a small value of $\bar{f} = 0.023$. We choose the drag coefficient of the medium $\tilde{\eta}_m$ so that the ratio between $\ell_{tissue}$, Eq. (18), and the size of the tissue is about 1. Therefore, we do not expect prominent wrinkling at the tissue scale. We choose $h_{cortex}/h_0 \approx 0.1$, a large value given the fact that $h_0$ is small in the very stretched initial state. We choose the cytosolic viscosity such that $\ell_{cortex}/R = 0.7$, Eq. (12), and therefore we expect cortical wrinkling.

In the uniaxial simulation in Movie 9, the parameters are the same except $\gamma_a^{lat}/\gamma_a^{ab} = 0.2$, $\bar{\gamma} = 0.5$, the ratio between $\ell_{tissue}$ and the size of the tissue is 0.7, and $\ell_{cortex}/R = 0.2$. Accordingly, we obtain a coexistence of tissue and cortical wrinkling and smaller cortical wrinkles.

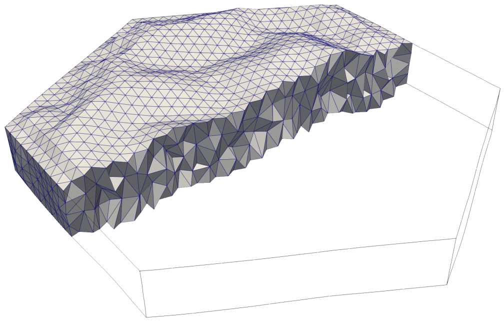

**Supplementary Figure 2:** Illustration of the bulk finite element mesh used in the computational model to model cytosolic viscosity. The boundary of this mesh is used to approximate the cortical surface using subdivision surface finite elements.

#### Supplementary References

[55] Salbreux, G., Prost, J. & Joanny, J.-F. Hydrodynamics of cellular cortical flows and the formation of contractile rings. *Physical Review Letters* **103**, 058102 (2009).

[56] Huang, R. & Suo, Z. Instability of a compressed elastic film on a viscous layer. *International Journal of Solids and Structures* **39**, 1791–1802 (2002).

[57] Huang, R. & Suo, Z. Wrinkling of a compressed elastic film on a viscous layer. *Journal of Applied Physics* **91**, 1135–1142 (2002).

[58] Huang, R. Kinetic wrinkling of an elastic film on a viscoelastic substrate. *Journal of the Mechanics and Physics of Solids* **53**, 63–89 (2005).

[59] Torres-Sánchez, A., Millán, D. & Arroyo, M. Modelling fluid deformable surfaces with an emphasis on biological interfaces. *Journal of fluid mechanics* **872**, 218–271 (2019).
